# A Conserved Landscape of Chemokine Receptor Co-expression Defines the Functional States of CD8+ T Cells in Melanoma

**DOI:** 10.64898/2025.12.10.693486

**Authors:** Rodney Macedo, David. W. Harle, Kevin Hoffer-Hawlik, Ximi K. Wang, Thomas McMahon-Skates, Alexandra Matschiner, Kirubel Belay, Yvonne M. Saenger, Benjamin Izar, Elham Azizi, Ran Reshef

## Abstract

Cancer immunotherapies, from checkpoint blockade to adoptive cell therapies like tumor-infiltrating lymphocytes (TILs), have revolutionized cancer treatment but are limited by variable efficacy and significant toxicities. A central challenge is identifying ideal T-cell populations that effectively eliminate tumors without causing off-target damage, a distinction not captured by existing biomarkers. We show that co-expression patterns of chemokine receptors (CRs) CXCR3, CCR5, and CXCR6 on CD8^+^ T cells provide a functional “code” defining subsets with divergent roles in on-target immunity versus off-target inflammation. In mouse and human melanoma, a triple-positive (CXCR3^+^CCR5^+^CXCR6^+^) T-cell subset is essential for tumor control, and its genetic signature correlates with positive clinical response, while a distinct CCR5^+^CXCR6^+^ subset drives liver immune-related adverse events (IRAEs). Crucially, this CR code reveals that immunotherapy actively reshapes T-cell trafficking patterns, uncovering profound heterogeneity within conventional populations and distinguishing potent anti-tumor progenitors from cells predisposed to exhaustion or off-target migration. This work establishes CR co-expression as a practical tool, providing a surface marker-based strategy to identify and enrich optimized T cells for adoptive therapies, thereby offering a framework to uncouple efficacy from toxicity.

**One Sentence Summary:** Co-expression of CCR5, CXCR6, and CXCR3 provides a functional ’code’ that separates T-cell-mediated anti-tumor efficacy from off-target toxicity, enabling the selection of superior cells for safer and more effective cancer immunotherapies.

**Graphical Abstract:** 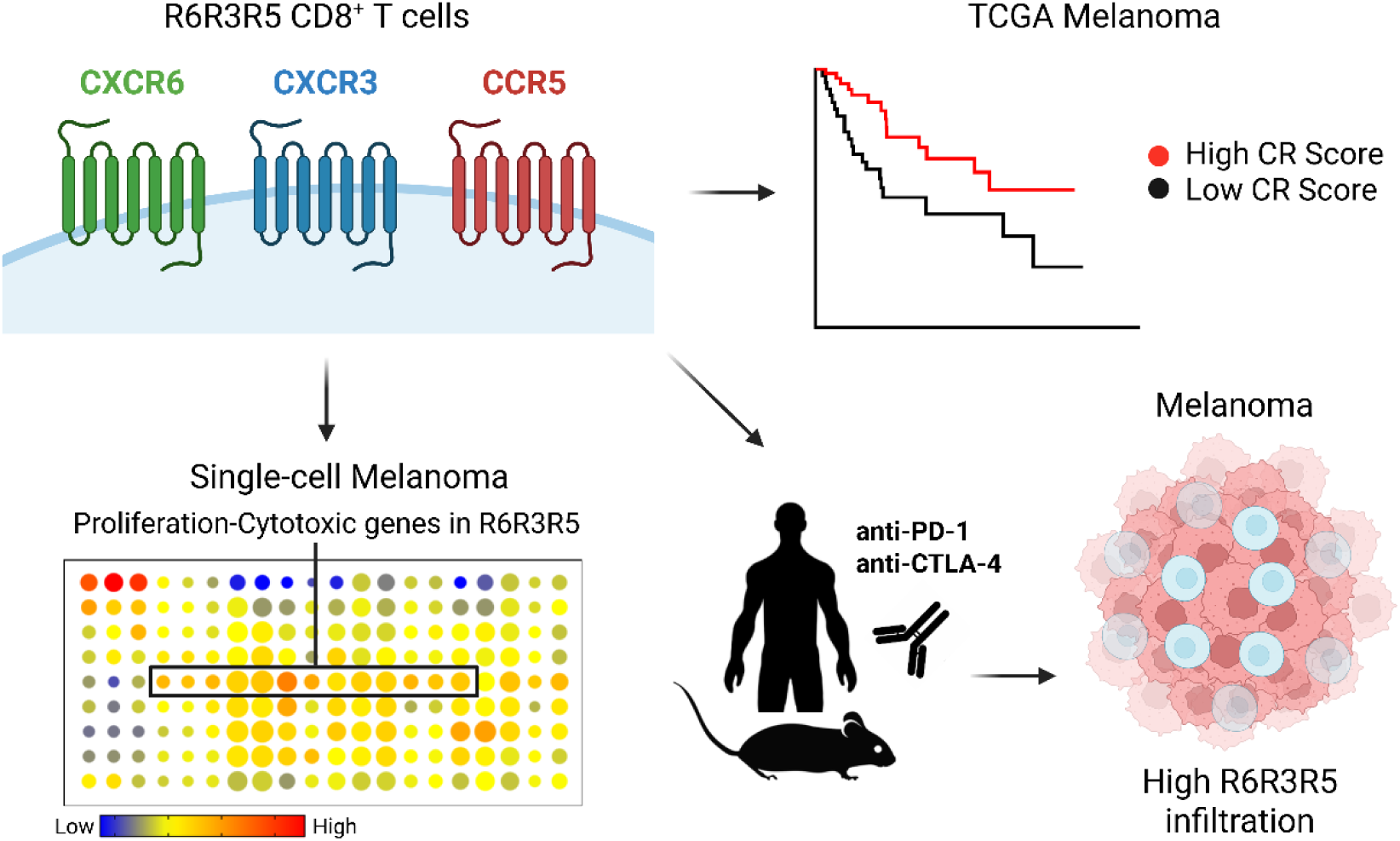

**Highlights:**

- · CXCR6, CXCR3 and CCR5 co-expression signature stratifies patient survival in human melanoma
- · CD8^+^ T cells co-expressing CXCR6, CXCR3 and CCR5 are critical effectors with high proliferative, cytotoxic, and activation profile in human and mice melanoma
- · CD8^+^ T cells co-expressing CXCR6, CXCR3 and CCR5 drive anti-tumoral responses during checkpoint blockade in both human and mice

---

Combined PD-1 and CTLA-4 blockade (P1C4) leads to impressive and durable clinical responses in melanoma ^1^, and many other cancers ^2,3^. Though transformative, only a subset of patients responds, and immune-related adverse events (IRAEs) account for the high incidence of grade 3-4 adverse events, treatment-related mortality and high discontinuation rates, limiting broader application of combination therapies ^4–11^. Both tumor rejection and IRAEs induced by checkpoint inhibitors are fundamentally governed by lymphocyte migration into tumors and healthy tissues ^12–14^. The recent approval of autologous tumor-infiltrating lymphocytes (TILs) in metastatic melanoma and the fact that the majority of patients do not achieve long-term benefit from this approach ^15–17^ further underscore the need for deeper understanding of T-cell populations with an ideal profile that combines effective trafficking to the tumor, potent effector function, high proliferative capacity (stemness), ability to resist suppressive signals, and low exhaustion state, resulting in an effective anti-tumor response.

Chemokines are proteins that control the trafficking of T cells and other leukocytes into different tissues by interacting with chemokine receptors (CR) expressed on the cell surface ^18^. Previous studies have investigated the role of CRs individually. Elevated expression of certain chemokine receptors such as CXCR3 and CCR5 or their ligands is associated with increased intratumoral T-cell infiltration and favorable outcomes in melanoma ^19–21^ and other cancers ^22–27^. These studies led to the experimental use of CR inhibitors in cancer patients with mixed results ^28–31^, possibly due to failure to account for the biologic reality that leukocytes co-express multiple CRs which collectively orchestrate their migration and potentially other functions ^32^. This highlights a significant gap in our understanding and an unmet need in identifying T-cell populations with an ideal profile -combining potent tumor trafficking and effector function while avoiding healthy tissues. This may facilitate the development of novel pharmacologic or adoptive cell therapy approaches in cancer.

Here, we address this challenge by investigating the co-expression patterns of key inflammatory CRs. We provide evidence in both mice and humans that the coordinated expression of CXCR3, CCR5, and CXCR6 defines distinct CD8^+^ T-cell subsets with divergent roles in mediating tumor clearance and immune-related toxicity following dual checkpoint blockade. Our findings may help design future therapies that modulate leukocyte migration to improve the efficacy and reduce immune-related toxicity of cancer immunotherapies.

## A T-cell Chemokine Receptor Signature Stratifies Patient Survival in Melanoma

To identify CRs central to the anti-tumor T-cell response and conserved across multiple cancers, we conducted an *in-silico* analysis of 11,323 tumor transcriptomes from 33 human cancers in the Cancer Genome Atlas (TCGA). This screen revealed that the inflammatory CR genes *CXCR3*, *CCR5*, and *CXCR6* exhibit both high expression and variance (Extended Data Fig. 1a). This receptor triad forms a distinct signature that strongly correlated with T-cell specific genes compared to other leukocyte populations (Fig. 1a top and Supplementary Table 1). This patter was conserved in 475 human melanoma samples (Fig. 1a bottom and Extended Data Fig. 1b). Consistent with this, the primary ligands for CXCR3, CCR5 and CXCR6 correlated with T-cell- and macrophage-specific genes, but not with tumor cells, B cells or fibroblasts (Extended Data Fig. 1c).

**Figure 1.**
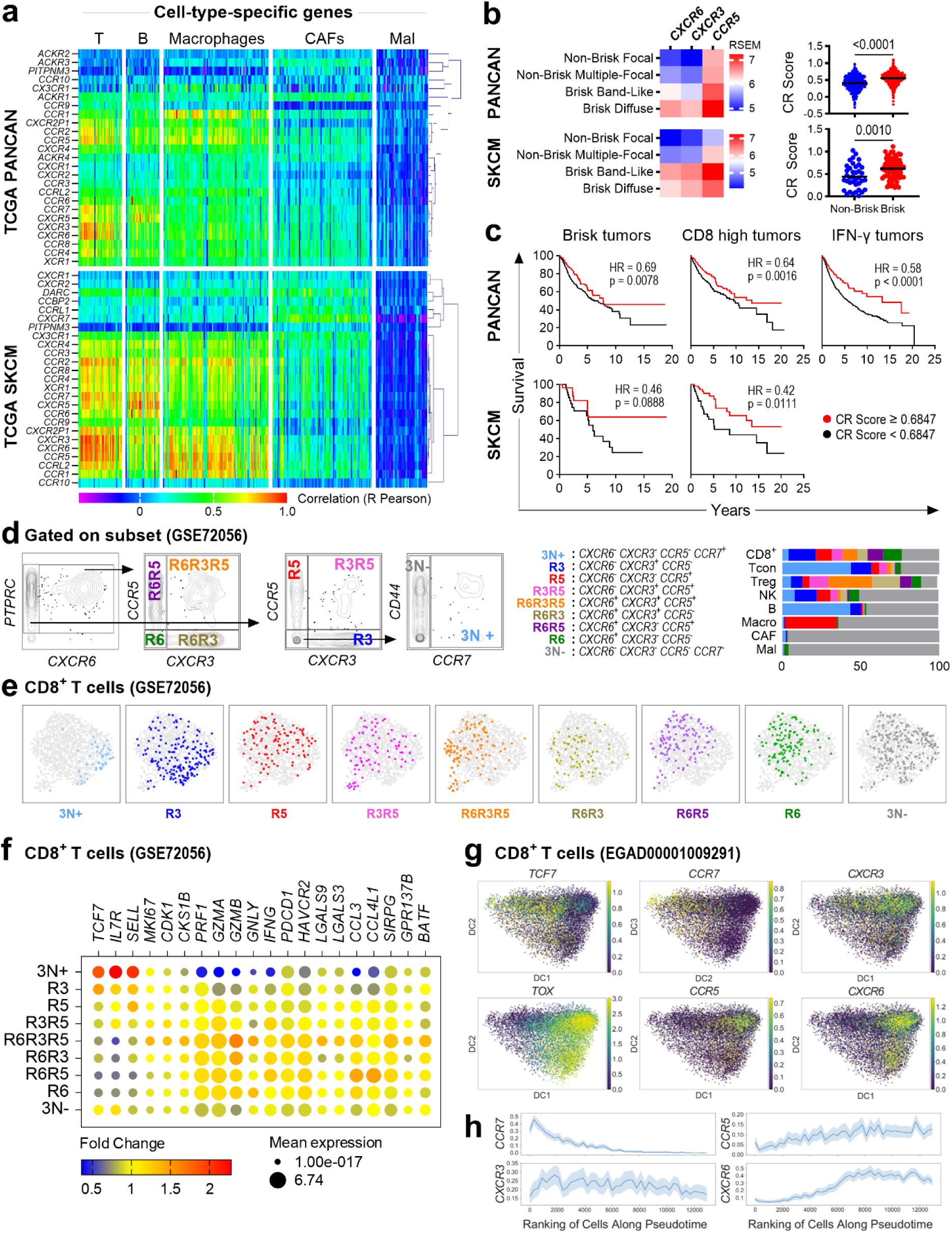
The landscape of chemokine receptor (CR) gene expression in human cancer defines functional T-cell subsets with prognostic importance. **a,b.** *In-silico* analysis of tumor transcriptomes from the TCGA Pan-Cancer Atlas (PANCAN; n=11323) and melanoma (SKCM; n=475) datasets. a. Hierarchical clustering heatmap showing the correlation of CR gene expression with cell-type-specific genes (Supplementary Table 1) in the tumor microenvironment. **b**. Heatmaps of *CXCR6*, *CXCR3* and *CCR5* expression (left) and CR scores (right) in PANCAN and melanoma samples, stratified by “brisk” lymphocytic infiltration status. **c**. Kaplan-Meier survival curves stratified by CR score in brisk, CD8-high or IFN-γ dominant tumors. **d-f**. Single-cell transcriptomic analysis of metastatic melanoma (GSE72056). **d**. Gating strategy defining CD8^+^ T-cell subsets based on *CXCR6*, *CXCR3*, *CCR5*, and *CCR7* co-expression (left) and heterogeneity of CR subsets across immune populations (right). **e**. UMAP projection showing CR-defined CD8+ T-cell subsets. **f**. Dot plot of gene expression for markers of naive, cytotoxicity, exhaustion, transcription factors and cytokines across CR-defined subsets. **g, h**. Diffusion mapping and pseudotime inference analysis from metastatic melanoma (EGAD00001009291) showing differential enrichment **(g)** and pseudotemporal expression patterns **(h)** of *CCR7*, *CXCR3*, *CCR5* and *CXCR6* along the *TCF7*-to-*TOX* differentiation trajectory.

While individual receptor expression was higher in “brisk” tumors ^33^ (characterized by dense lymphocytic infiltrate) versus “non-brisk” PANCAN and melanoma tumors, a combinatorial CR score proved superior for stratification. Specifically, CR scores (based on *CXCR6*, *CXCR3* and *CCR5* expression) were not only associated with significant differences between these tumor types (Fig. 1b, Extended Data Fig. 1d) but also linked to improved overall survival uniquely within brisk tumors (Fig. 1c). Furthermore, CR scores correlated with higher abundance of CD8^+^ T cells ^34^ and IFN-γ signatures ^35^ in tumors, predicting a survival benefit even within CD8^+^ and IFN-γ-dominant tumors (Fig. 1c, Extended Data Fig. 1e-g). This was reinforced by an analysis of transcriptional data from metastatic ^36^ and primary ^37^ human melanoma confirming that higher CR scores associate with non-recurrent (Extended Data Fig. 1h) and responder (Extended Data Fig. 1i) tumors, respectively. Differentially expressed genes (DEGs) in samples with high *CXCR6*, *CXCR3*, and *CCR5* co-expression were enriched for T cell activation pathways (Extended Data Fig. 1j and 1k). Collectively, these data establish *CXCR3*, *CCR5*, and *CXCR6* as a core inflammatory T-cell signature associated with improved survival.

## Combinatorial Chemokine Receptor Expression Reveals a Hierarchy of T-cell Effector Programs

To determine if CR co-expression patterns define distinct functional states, we performed *in-silico* single-cell analysis of treated and untreated human metastatic melanoma tumors ^38^. Focusing on CD8^+^ T cells (CD8^+^), after identifying major cell populations (Extended Data Fig. 2a) using cell type-specific genes, we classified nine unique subsets based on the combinatorial expression of *CXCR6*, *CXCR3* and *CCR5* (using *CCR7* to demarcate naive cells) (Fig. 1d left). This approach revealed extensive CR heterogeneity in the tumor microenvironment (TME) (Fig. 1d right) and demonstrated that multi-receptor patterns, rather than single markers, are required to delineate functionally distinct cell states.

Transcriptional profiling of CR-defined subsets uncovered a functional continuum across CD8^+^ T cells, from naive to terminally differentiated states (Fig. 1e-f, Extended Data Fig. 2b-c, and Supplementary Table 2). At one end of the spectrum, the triple-negative (3N+) and CXCR3-only (R3) subsets were the least differentiated, expressing markers of stemness and a naive state, alongside chemokines involved in dendritic cells recruitment, suggesting a progenitor-like role. In contrast, co-expression of CCR5 and/or CXCR6 marked cells with potent effector programs. The **triple-positive (R6R3R5) subset** emerged as a critical effector phenotype, exhibiting the highest expression of proliferation, cytotoxicity, and activation genes. Dual-expressing subsets represented distinct intermediate states; R3R5 TILs were highly proliferative, while R6R3 TILs displayed a resident-effector profile with high expression of cytotoxic molecules but lower proliferation. Finally, CXCR3-negative TILs appeared terminally differentiated, characterized by high expression of exhaustion markers. Within this group, R5 cells resembled an immature memory state, while R6R5 and R6 subsets were more akin to short-lived effector cells, with R6R5 being more cytotoxic and R6 showing more tissue residency markers. These findings indicate that specific combinations of CXCR3, CCR5, and CXCR6 define a detailed functional map of CD8^+^ T cell diversity within the TME.

## Chemokine Receptor Expression Marks a Differentiation and Activation Trajectory in TILs

To map the evolution of CR expression during T cell maturation, we analyzed a second human metastatic melanoma single-cell RNA-seq dataset ^39^. We applied pseudotime inference (Extended Data Fig. 2d-e), a computational approach which orders cells along a continuous phenotypic trajectory, in this case from naive (*TCF7*-expressing) to terminally differentiated (*TOX*-expressing) CD8^+^ T cells ^40^. Sliding window analysis of gene expression revealed a distinct, sequential pattern of CR expression along this differentiation trajectory (Fig. 1g). The trajectory begins with high *CCR7* expression, which is progressively lost (Fig. 1h). *CXCR3* emerges early and is sustained, whereas *CCR5* appears and peaks in intermediate states. Finally, *CXCR6* expression is acquired late, marking the most differentiated cells and correlating with *TOX* expression. Taken together, these results suggest that T-cell maturation within the tumor involves programmed, sequential co-expression of *CXCR3*, *CCR5* and *CXCR6* followed by a gradual loss of these receptors in the terminally differentiated phase. This evolution underscores that dynamic combinatorial patterns, rather than static expression of a single receptor, are critical for accurately characterizing the functional state of TIL subsets and their distinct roles in mediating anti-tumor efficacy.

## A Conserved Chemokine Receptor Code Defines Functional T-cell Programs in Mice and Humans

We next validated these CR-defined subsets at the protein level using multiparametric flow cytometry in healthy human and mouse lymphocytes (Fig. 2a, Extended Data Fig. 3 and Supplementary Table 3). This analysis revealed that the CR co-expression “code” is a conserved feature that defines a functional hierarchy (Fig. 2b-c, Extended Data Fig. 4a-c). In both species, the least differentiated, naive-like subsets (3N+ and R3) were the most abundant and largely corresponded to traditional Naive (T_N_) and Central Memory (T_CM_) populations. In contrast, the Effector Memory (T_EM_) population was highly heterogeneous, comprising the full spectrum of CR-defined subsets (Fig. 2d-e, Extended Data Fig. 4d-f).

**Figure 2.**
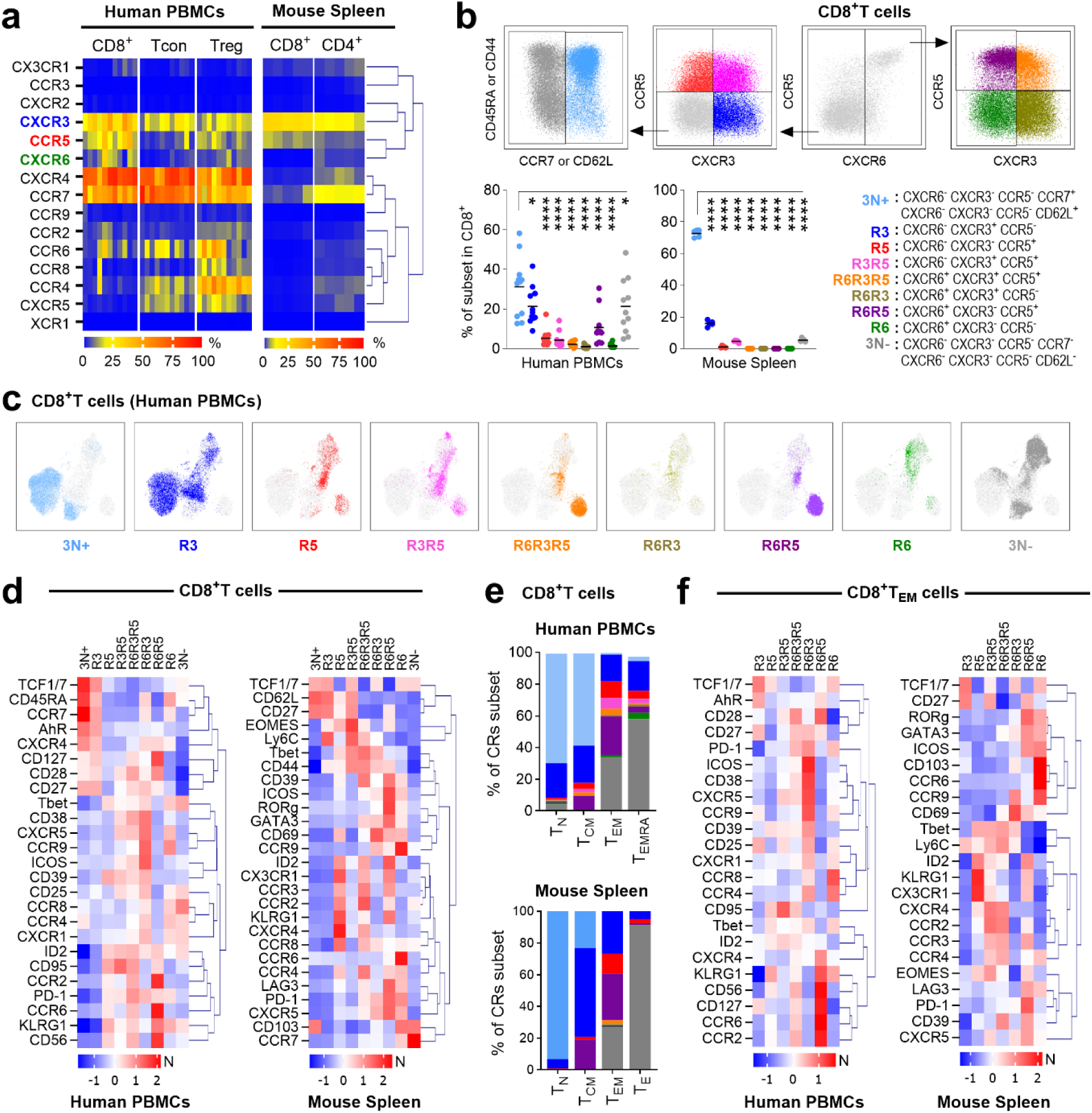
Co-expression of CXCR3, CCR5 and CXCR6 defines conserved functional T-cell subsets in humans and mice. **a.** Heatmap of CR co-expression in T-cell subsets from human PBMCs (n=11) and mouse splenocytes (n=5). **b.** Flow cytometry gating strategy identifying CR subsets. Scatter plots show the percentages of subsets in human (left) and mouse (right) CD8^+^ T-cells. **c**. UMAP projection of CR-defined subsets in human CD8^+^ T cells. **d**. Heatmap of normalized (N) protein expression of CRs, transcription factors, activation and exhaustion markers in human and mouse CD8^+^ T cells. **e**. Distribution of CR-defined subsets within Naive (T_N_), Central Memory (T_CM_), Effector Memory (T_EM_) and effector (T_E_ or T_EMRA_) populations in human and mice. **f**. Hierarchical clustering heatmap of normalized (N) protein expression of CRs, transcription factors, activation and exhaustion markers specifically in human and murine CD8^+^ T_EM_ populations. Statistical significance determined by unpaired Mann-Whitney tests, *P < 0.05, **P < 0.01, ***P < 0.001, ****P < 0.0001.

Crucially, our CR framework uncovered profound functional divergence within the classically defined TEM population (Fig. 2f and Extended Data Fig. 4g). For example, R3 T_EM_ cells retained stem-like properties (high TCF1/7), while subsets co-expressing multiple CRs, such as R6R3R5, were primed for cytotoxicity and proliferation. Other subsets showed exhausted phenotypes (high PD-1, LAG3) or distinct tissue-homing potentials, expressing receptors that support either tumor infiltration (CCR2) or mucosal sites (CCR6, CCR9). These results confirm at the protein level that CR co-expression defines conserved, functionally distinct T-cell subsets and resolves critical heterogeneity within the traditional T_EM_ population, distinguishing cells with potent anti-tumor potential (R6R3R5) from those predisposed to exhaustion or migration to other tissues.

## Distinct T-cell CR Subsets Drive On-Target Efficacy Versus Off-Target Inflammation During Checkpoint Blockade

To dissect the *in vivo* roles of CR-defined subsets in the context of immunotherapy, we studied their response to dual PD-1 and CTLA-4 blockade (P1C4) in a B16-F10 melanoma model (Fig. 3a). This combination was chosen as it was most effective at increasing CD8^+^ T_EM_ cells compared to single-agent therapy (Extended Data Fig. 5a) and is clinically relevant due to its high efficacy in human melanoma. Treatment increased CD8^+^ T and T_EM_ percentages in both tumor and liver (Fig. 3b). While liver enzymes remained unchanged in this model, immunohistochemistry revealed a significant increase in CD3^+^ T cells in the liver, indicating subclinical inflammation (Extended Data Fig. 5c-e). These results established a robust model to simultaneously investigate tumor control and tissue-specific drivers of early immune-related toxicity.

**Figure 3.**
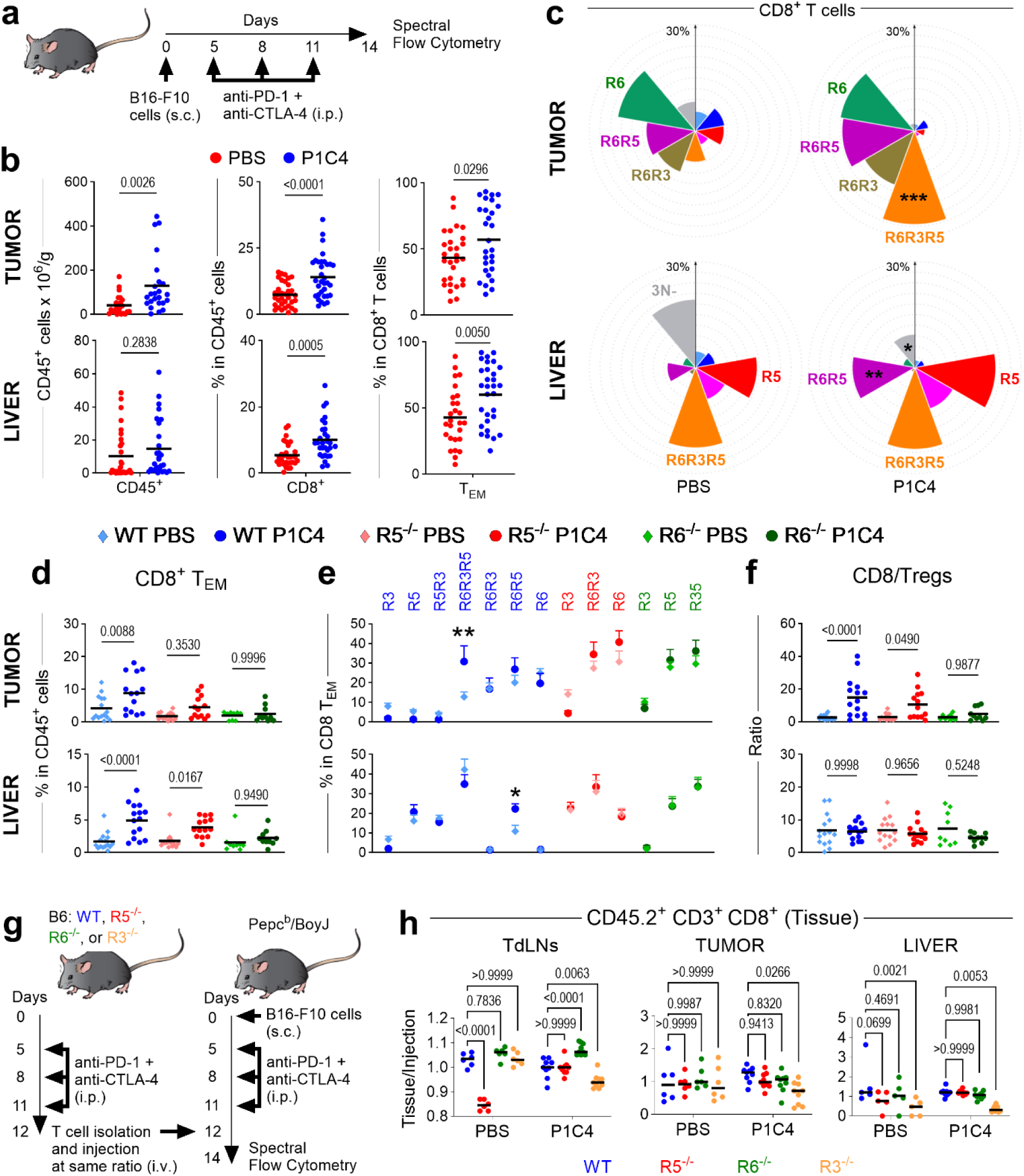
Triple expression of CXCR3, CCR5 and CXCR6 is required for PD-1/CTLA4-induced effector T-cell increase in melanoma. **a**. Experimental schematic: B16-F10 tumor-bearing C57BL/6J mice (n=15) were treated with PD-1/CTLA-4 (P1C4) or PBS. Cells from tumors and liver were isolated on day 14 and analyzed by spectral flow cytometry. **b.** Quantification of tumor-infiltrating CD45^+^, CD8^+^, and CD8^+^ T_EM_ cells in tumor and liver. **c**. Nightingale Rose charts showing the frequency of CR-defined subsets in tumor and liver after P1C4 treatment (* to *** representing significant differences between P1C4 and PBS). **d-f**. Impact of *Ccr5* (R5) or *Cxcr6* (R6) deficiency on tumor (top) and liver (bottom). **d**. CD8^+^ T_EM_ infiltration in WT, *Ccr5*^-/-^ or *Cxcr6*^-/-^ mice. **e**. Frequency of R6R3R5 and R6R5 subsets. **f.** CD8^+^/Treg ratio. **g.** Competitive homing assay schematic. **h**. Homing ratios of CD45.2^+^ CD3^+^ cells in TdLNs, tumor and liver. Statistical significance determined by unpaired ANOVA or Mann-Whitney multiple comparisons tests, *P < 0.05, **P < 0.01, ***P < 0.001, ****P < 0.0001.

P1C4 therapy profoundly and selectively altered the distribution of CR-defined subsets in peripheral tissues (Fig. 3c). In the tumor, treatment drove a dramatic expansion of the triple-positive R6R3R5 subset. Conversely, in the liver, P1C4 selectively increased the R6R5 subset. These tissue-specific effects were not observed in the spleen or tumor-draining lymph nodes (TdLNs)(Extended Data Fig. 5f). This divergence suggests that distinct CR programs mediate on-target anti-tumor activity versus off-target immune infiltration. This conclusion is also supported by a corresponding increase in the ligands for CXCR3, CCR5 (e.g., CXCL9, CXCL10, CCL5) in both tumor and liver supernatants after treatment, whereas CXCR6 ligand (CXCL16) is higher only in the tumor (Extended Data Fig. 5g).

Using genetic knockout models, we demonstrated that CR co-expression is essential for therapeutic efficacy. In *CCR5*- or *CXCR6*-deficient mice, P1C4 failed to increase intratumoral CD8^+^ T_EM_ cells, losing any significant change in CR-defined subsets (Fig. 3d-e). Furthermore, the P1C4-mediated improvement in the CD8^+^/Treg ratio was blunted in *CCR5*-knockout mice and eliminated in *CXCR6-*knockout mice (Fig. 3f). These data demonstrate that an optimal anti-tumor response to P1C4 requires the functional contribution of T cells able to express all three receptors.

A competitive short-term homing assay further clarified CR roles (Fig. 3g). As expected, CXCR3 is critical for T-cell trafficking into the tumor, TdLNs, and liver following P1C4 treatment (Fig. 3h and Extended Data Fig. 5h-i). The roles of the other receptors are more complex; for instance, *CXCR6*-deficient cells accumulate in lymphoid tissues after P1C4 treatment, suggesting impaired migration to the tumor or inability to establish tissue residency status. Altogether, these data establish that the functional outcome of P1C4 therapy—tumor rejection versus immune-mediated toxicity—relies on the specific co-expression of CXCR3, CCR5, and CXCR6 on responding T cells.

## Anti-Tumor T-cell Subsets Defined by CR Co-expression Expand with Checkpoint Blockade and Correlate with Clinical Response

To validate our findings clinically, we analyzed PBMCs from melanoma patients, comparing Immune Checkpoint Blockade (ICB)-treated and ICB-naive patients (Supplementary Table 4). ICB-treated patients had significantly higher percentages of CD8^+^ T cells in their blood compared to ICB-naive patients (Extended Data Fig. 6a-b). Strikingly, dual ICB treatment led to a significant expansion of the circulating triple positive R6R3R5 subset (Fig 4a), mirroring our findings in murine splenic CD8^+^ T cells (Extended Data Fig. 6c).

**Figure 4.**
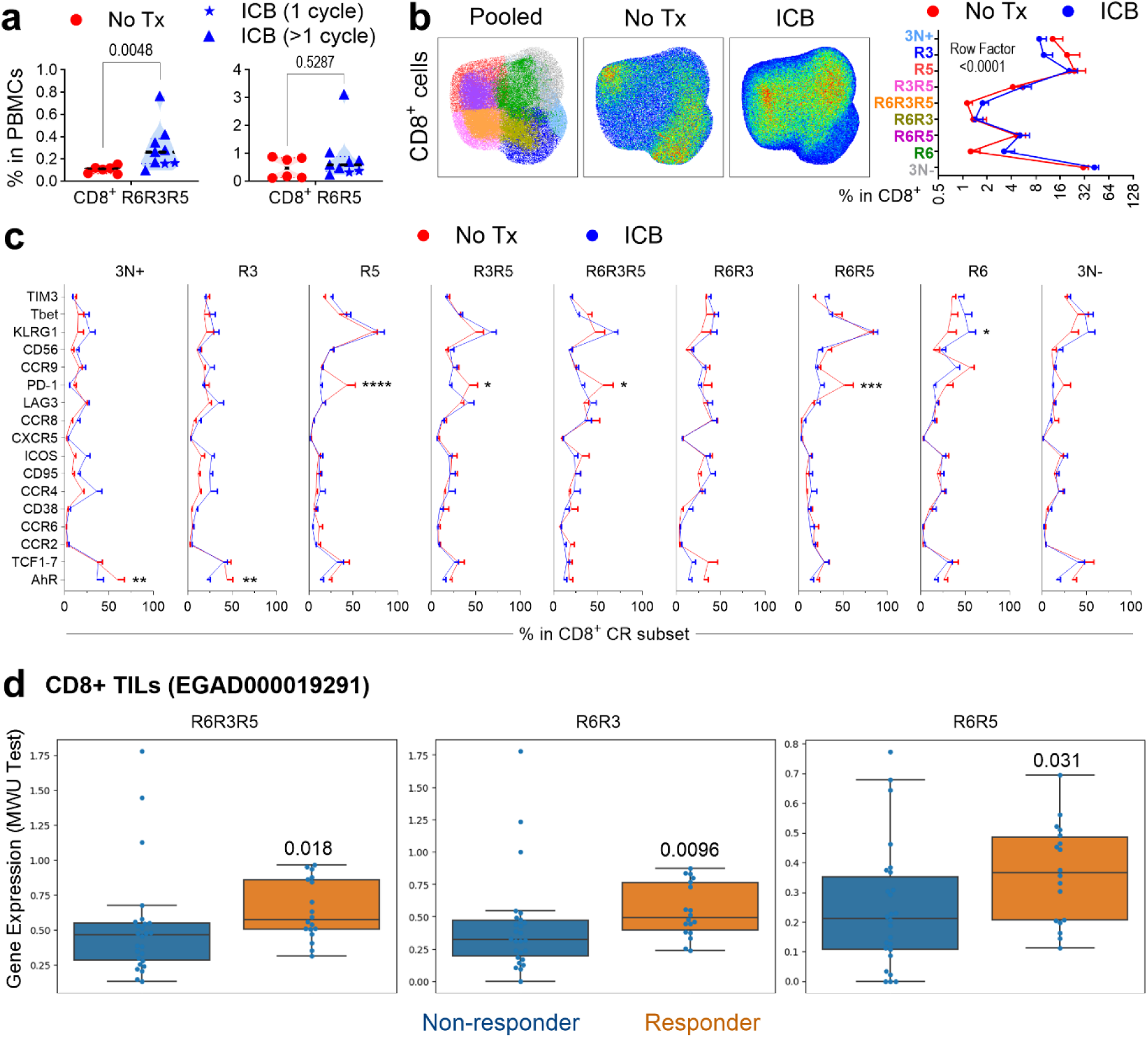
PD-1/CTLA4 blockade reshapes CR co-expression in peripheral T cells of melanoma patients. **a**. Frequency of circulating CD8^+^ R6R3R5 and R6R5 in ICB-treated patients (n = 9) versus untreated patients (No Tx, n = 6). **b**. UMAP projection (left) and profile plots (right) showing shifts in CR subset distribution of CD8^+^ T cells following ICB. **c**. Profile plots showing phenotypic changes in CRs, transcription factors, and functional markers within CD8^+^ CR-defined subsets after ICB. **d**. Expression of R6R3R5, R6R3, and R6R5 gene signatures in ICB responders versus non-responders (EGAD000019291). Statistical significance determined by unpaired ANOVA or Mann-Whitney multiple comparisons tests, *P < 0.05, **P < 0.01, ***P < 0.001, ****P < 0.0001.

ICB remodeled the overall profile and immunophenotype of CR-defined subsets (Fig. 4b-c). Interestingly, ICB treatment was associated with significantly decreased AhR in 3N+ and R3 subsets, while R6 subsets exhibited increased KLRG1. Several subsets (R5, R3R5, R6R3R5, R6R5) showed declined PD-1 expression, suggesting receptor occupancy by therapeutic antibody ^41^. These results confirm in humans that P1C4 treatment increases the proportion of CR-defined subsets responsible for tumor control but also modifies the functional properties of these subsets.

Finally, to determine if CR-defined subsets within the human TME are associated with clinical outcomes, we analyzed the single-cell transcriptional dataset from metastatic melanoma patients treated with ICB ^39^. This analysis revealed that *CXCR6* expression was significantly higher (*P*=0.0095) in responder than non-responders (Extended Data Fig. 6d). Importantly, gene signatures for CXCR6-co-expressing subsets -R6R3R5, R6R3 and R6R5- were significantly enriched in tumors of responder patients (*P*<0.05) (Fig. 4d). These results confirm that ICB reshapes the circulating T cell landscape to favor the highly functional R6R3R5 subset. Critically, they also establish that the enrichment of CXCR6-co-expressing T-cell subsets within the tumor itself is a key feature of a successful anti-tumor response to immunotherapy.

## Discussion

A central challenge in cancer immunotherapy—from checkpoint blockade to adoptive cell therapies like TILs—is uncoupling potent anti-tumor efficacy from treatment-limiting, off-target toxicity. This requires a deeper understanding of the T-cell subsets that mediate these opposing outcomes. While the roles of individual chemokine receptors in T-cell trafficking are well-studied, the combinatorial “code” they form to dictate specific functional states has remained unclear. Our study addresses this gap, demonstrating that the co-expression of CXCR3, CCR5, and CXCR6 provides a conserved functional map of CD8^+^ T cells in both mice and humans, allowing for the prospective identification of T-cell subsets responsible for on-target efficacy versus off-target inflammation.

Our analysis reveals that this CR code defines a fundamental and conserved T-cell developmental trajectory. In healthy individuals, we found that the hierarchy progresses from naive-like, progenitor cells expressing primarily CXCR3 (R3 subset) to highly differentiated effector cells co-expressing all three receptors (R6R3R5 subset). Crucially, this framework uncovers profound functional heterogeneity within traditionally defined populations like effector memory T cells (T_EM_). Within the T_EM_ pool, our code distinguishes stem-like progenitors (R3) from highly cytotoxic and proliferative effectors (R6R3R5) and exhausted subsets, a level of functional resolution not achievable with conventional markers. Importantly, our analysis demonstrates that specific combinations of these three receptors, rather than single markers, are required to accurately define the functional state and potential of a T cell.

In the context of cancer immunotherapy, this framework provides a powerful lens to dissect the mechanisms of both tumor response and IRAEs. Our preclinical data show that dual ICB selectively expands the triple-positive R6R3R5 subset in the tumor, a population essential for therapeutic efficacy. Conversely, the therapy promoted the accumulation of a distinct CCR5^+^CXCR6^+^ (R6R5) subset in the liver, a common site of IRAEs. This finding mechanistically links a specific, identifiable T-cell subset to off-target toxicity, providing a cellular target for future strategies aimed at mitigating IRAEs. The fact that genetic deletion of either CCR5 or CXCR6 abrogated the therapeutic effect of ICB underscores that the coordinated function of all three receptors on the R6R3R5 subset is non-redundant and essential for a successful anti-tumor response.

Ultimately, our findings set up this CR code not just as a prognostic biomarker but as a practical tool for developing better immunotherapies. The prognostic value of single chemokine receptors has been inconsistent across studies, likely because, as we show, the functional output of T cells is associated with receptor combinations. The true translational potential of our work lies in its direct application to cell-based therapies. The surface markers of this code are readily detectable by high-dimensional flow cytometry, offering a straightforward strategy to identify and prospectively isolate the most desirable T-cell populations. For adoptive cell therapies, such as TILs, this framework provides a clear rationale for enriching the final product with the highly proliferative and cytotoxic R6R3R5 subset while potentially depleting populations like R6R5 that may contribute to toxicity. This approach offers a tangible path forward to engineer more potent and safer immunotherapies, improving the therapeutic window by maximizing anti-tumor activity while minimizing collateral damage.

While the expression profile of CRs and other trafficking molecules in immune cells has been previously described in healthy humans ^42,43^ and in cancer ^44,45^, the functional impact of their co-expression on the anti-tumor response requires multiparametric, single-cell analysis. Using genetic datasets and high-dimensional flow cytometry, our study simultaneously analyzed multiple CRs with functional markers and transcription factors to clarify the critical role of CR co-expression in shaping anti-tumor T-cell responses.

## Data and code availability

All data generated in this study are included in this published article and its supplementary information. Code relating to data processing and figure generation is available in our GitHub repository. Patient information is available on the supplementary table 4. Any additional information required to reanalyze the data reported in this paper is available from the corresponding authors upon request.

## Acknowledgements

We thank the CCTI Biobank and Flow Cytometry Cores. We gratefully acknowledge the Columbia Institutional Review Board and all patients who generously consented to participate in this study, as well as the Institutional Animal Care and Use Committee. We are also grateful to Megan Sykes, Siu-Hong Ho, Audrey Baeyens, and Claudia Gutierrez-Chavez for helpful discussions.

## Author contributions

R.M., D.H., and R.R., conceived the study. R.M., D.H., K.H.H., X.W., A.M., and K.B. performed sample processing and data collection. R.M., D.H., K.H.H., X.W., T.M.S., K.B., Y.S., B.I., E.A., and R.R. provided clinical samples and data and maintained regulatory oversight. R.M., D.H., K.H.H., X.W., and T.M.S., analyzed the data. R.M., D.H., K.H.H., X.W., T.M.S., A.M., K.B., Y.S., B.I., E.A., and R.R. interpreted the data. R.M., R.R., E.A. and K.H.H. wrote the manuscript.

## Methods

### Human samples

All study participants signed a written informed consent before inclusion in the study, and the protocols were approved by the Columbia University institutional review board (IRB). Peripheral blood mononuclear cells (PBMCs) were obtained from melanoma patients by Ficoll gradients and cryopreserved. PBMCs from healthy donors were obtained by Ficoll gradients and freshly analyzed using spectral flow cytometry. Characteristics of patients and healthy donors are summarized in Supplementary Table 4.

### Cell lines

B16-F10 cell line was obtained from ATCC. B16-F10 cells were cultured at 37°C in Dulbecco’s Modified Eagle Media, supplemented with 10% FBS, 100 U ml^-1^ penicillin and 100 μg ml^-1^ streptomycin, 2 mM L-glutamine. All cell lines were determined to be free of Mycoplasma (Lonza) and common mouse pathogens (IDEXX).

### *In vivo* mouse studies

All mice were used according to approved protocols by Columbia University Institutional Animal Care and Use Committee (IACUC). Female mice, including C57BL/6J (B6) (Jax #000664), B6.129P2-*Ccr5^tm1Kuz^*/J (Jax #005427), B6.129P2-*Cxcr6^tm1Litt^*/J (Jax #005693), B6.129P2-*Cxcr3^tm1Dgen^*/J (Jax #005796), and B6.SJL-*Ptprc^a^Pepc^b^*/BoyJ (Jax #002014) were obtained from The Jackson Laboratory at 7-12 weeks of age and were used for studies one week or more after their arrival in the animal barrier facility of the Columbia Center for Translational Immunology. For B16-F10 melanoma, 1 × 10^5^ B16-F10 cells diluted in PBS were mixed with an equal volume of Matrigel (Corning) and subcutaneously injected in the right flank of B6 mice on day 0. Blocking antibodies were given on days 5, 8 and 11 after tumor implantation. Antibodies used for *in vivo* immune checkpoint blockade experiments were given intraperitoneally at a dose of 200 mg per mouse and included: CTLA4 (9H10) and PD-1 (RMP1-14) (BioXCell). Tumor diameters were measured using calipers. Volume was calculated using the formula: (length × width^2^)/2. Mice were euthanized when tumor burden reached a volume of 1500 mm³. Normalized tumor response to treatment is the measured volume at time t (Vt) relative to volume at baseline (Vo).

### Preparation of murine single-cell suspensions for flow cytometry

Murine cell suspensions were obtained from tumors or livers by a combination of GentleMACS (Miltenyi Biotec) dissociation and enzymatic dissociation using 1 mg/mL of collagenase IV (Sigma-Aldrich) and 0.2 mg/mL of DNase (Roche), followed by incubation with ACK Lysis buffer (Life Technologies) for 1 minute to remove red blood cells. For tumor-draining lymph nodes, mechanical and enzymatic dissociation was performed. After enzymatic incubation, samples were filtered through a 70-µm nylon cell strainer (Falcon). Splenocytes were obtained by dissociation using a 70-µm nylon cell strainer (Falcon) followed by ACK lysis buffer incubation for 1 minute. Cell suspensions were counted using Trypan Blue for cell viability.

### Flow cytometry and cell sorting

For flow cytometric analysis, live/dead cell discrimination was performed using Zombie NIR Fixable Viability dye (Biolegend) or eFluor780 Fixable Viability Dye (eBiosciences). Fc receptors were blocked using 2.4G2 anti-mouse CD16/CD32 (BD Pharmingen) or Human TruStain FcX (Biolegend). Cell surface staining was done for 15 min at 36°C, followed by 15 min at 4°C. Intracellular staining was done using a fixation/permeabilization kit (eBioscience). For human cells, we used multiparametric spectral flow cytometry with a 43-color extracellular panel and 42-color intracellular panel that allowed us to study multiple leukocyte populations, CRs, and functional markers simultaneously. In a similar way, for murine cells we used a 36-color extracellular panel and a 35-color intracellular panel to study multiple leukocyte populations, CRs, and other markers. For certain murine experiments traditional flow cytometry was used to identify T cell populations. All flow cytometric analyses were performed using Aurora 5 lasers (Cytek) or Fortessa II (BD) and analyzed using FlowJo software (BD). Cell sorting of CR-defined subsets was performed using an Influx Cell Sorter (BD). Human and murine sorted cells were stained with CellTrace kit followed by stimulation with Phorbol 12-myristate 13-acetate (PMA) for 2 hours to study activation or 2 days for proliferation. See Supplementary Table 3 for a list of antibodies used.

### Competitive homing experiment

To determine the role of each individual CR in homing and tissue-infiltration of T cells, we performed a competitive short-term homing strategy ^47^ to identify four groups of cells injected at identical proportions. First, we treated WT, CCR5^-/-^, CXCR6^-/-^, or CXCR3 -/- mice with anti-PD-1 + anti-CTLA-4 (P1C4) or PBS on days 5, 8, and 11. On Day 12, mice were euthanized, and T cells were isolated from the spleen using Pan T Cell Isolation Kit II (Miltenyi Biotec). After labelling T cells with different CellTrace Kits (except R6^-/-^ mice which express GFP reporter) and combining them at similar proportion, 15 × 10^6^ T cells were injected into the tail vein of B16 tumor-bearing PepBoy mice (Day 12 after tumor injection), also treated with PBS or P1C4 following the same treatment schedule and respecting the same group (PBS in PBS, P1C4 in P1C4). After 48 hours recipient mice were intravenously injected with anti-CD90.2 antibody for discrimination between vascular (circulating) and tissue lymphocytes ^48^. After 3 minutes, mice were sacrificed, and the percentage of WT, CCR5^-/-^, CXCR6^-/-^, or CXCR3^-/-^ in CD45.2^+^ CD3^+^ cells in each tissue was divided by the percentage of the same cells before injection.

### Immunohistochemistry of mouse tissues

For liver immunohistochemistry, rabbit anti-CD3ε antibody (Invitrogen SP7) was diluted 1:100 and incubated at room temperature for 2 hours. Signal amplification was performed using ImmPRESS Horse anti-rabbit IgG antibody-HRP (Vector) and DAB substrate Kit (BD Pharmingen) following the manufacturer’s protocol. Slides were counterstained and mounted.

### TCGA RNAseq

RNAseq data was downloaded from the TCGA Pan-Cancer and TCGA Melanoma (SKCM) cohorts from xenabrowser.net ^51^ (IDs: EB++AdjustPANCAN_IlluminaHiSeq_RNASeqV2.geneExp.xena and TCGA.SKCM.sampleMap/HiSeqV2; unit: log2(norm_count + 1)), and corresponding phenotype datasets (IDs: Survival_SupplementalTable_S1_20171025_xena_sp, survival/SKCM_survival.txt, TCGA_phenotype_denseDataOnlyDownload.tsv, TCGA.SKCM.sampleMap/SKCM_clinicalMatrix). This dataset contained 11060 samples for PANCAN and 464 samples for SKCM, but only primary and metastatic tumors were analyzed (n=10272 for PANCAN, n=104 for primary SKCM, and n=368 for metastatic SKCM). R value from Pearson correlation between normalized RSEM counts (> 5 log2(x+1)) of CRs/ligands and cell-type-specific genes (Supplementary Table 1; based on Tirosh et al) was calculated using GraphPad Prism v10. Then, an unsupervised hierarchical clustering gene tree was generated based on these correlations using Multi Experiment Viewer (MeV) v4.9 ^52^. For combinatorial CR scores, *CXCR6*, *CXCR3* and *CCR5* normalized gene expression were averaged (Score 1), or *CCR5* normalized gene expression was subtracted from *CXCR6* and *CXCR3* normalized gene expression sum (Score 2). For survival analysis, optimal cutoffs were calculated using Kaplan-Meier Plotter (kmplot.com) ^53^, and Kaplan-Meier curves and Log-rank p values were generated using GraphPad Prism v10. Identification of upregulated differentially expressed genes from TCGA and Liu *et al* primary melanoma samples was done using SeqGeq (BD), a platform that allows visualization of scRNA seq data and RNA seq data. STRING database (https://string-db.org/), was used to generate biological process gene ontology analysis ^54^.

### Analysis of publicly available single-cell datasets

To perform an *in-silico* analysis of single-cell RNA seq data from human metastatic melanoma, the GSE72056_melanoma_single_cell_revised_v2.txt.gz file was downloaded from GEO Series GSE72056. Using SeqGeq (BD), a characterization of immune and non-immune cells in the TME was performed with UMAP multidimensional analysis. Subsets were defined in CD8^+^ T cells by the co-expression of *CCR7*, *CXCR6*, *CXCR3* and *CCR5* following the gating strategy defined in Fig. 1d left. Finally, gene expression of other functional markers was measured within CR-defined subsets.

### scRNA-seq analysis of external clinical melanoma cohort immune cells

Pre-processed scRNA-seq datasets of immune cells (n=49,009) from patients studied in Pozniak et. al ^39^ were loaded into Seurat in R and subsequently Scanpy in Python. Immune cell annotation was guided by automatic annotation using SingleR and its built-in reference dataset of stroma and immune cells ‘BlueprintEncodeData’ as previously described ^49^. Non-cycling CD8^+^ T cell subtypes (n=13,304) were selected for downstream analysis in Scanpy. The top 8,000 highly variable genes were calculated using the ‘Seurat_V3’ method implemented in Scanpy. Gene expression counts were normalized to median total counts and logarithmized in Scanpy. CD8^+^ T cells from individual samples were batch corrected using an scVI model trained with two hidden layers and 30-dimensional latent space ^50^. After performing neighborhood graph construction using 20 nearest neighbors in scVI latent space, leiden clustering with default resolution, and evaluation of known marker genes and CD8^+^ T cell gene signatures in Scanpy, a noisy outlier cluster was removed to prevent distortions in subsequent diffusion maps. Then, after reconstructing the neighborhood graph using 300 nearest neighbors in scVI space, we computed diffusion components (DCs) using the Scanpy implementation to examine continuous phenotypic dynamics. The first and second DCs reflected expected *TCF7*^+^ to *TOX*^+^ progression of CD8^+^ T cells, and diffusion pseudotime along this trajectory was computed using the Scanpy implementation to reflect this progression. We then plotted the expression of *CCR7*, *CXCR3*, *CCR5*, and *CXCR6* across pseudotime using CD8^+^ T cells ordered by diffusion pseudotime rank. Given the sparsity of CR expression, which is typical for a clinical scRNA-seq dataset, we used sliding windows of 300 cells to average normalized expression across the CD8^+^ T cell trajectory. We also computed the 95% confidence interval on the same window of 300 cells to reflect variation in expression across the trajectory. The window size of 300 reduced sparsity while still conserving the expected negative binomial distribution of CR gene expression counts. To compare CR expression patterns over CD8^+^ T cell trajectory, expression of each CR was scaled and plotted across cell pseudotime rank, reflecting the expected progression of *CCR7*^+^ to *CXCR6*^+^ expression in CD8^+^ T cells. Finally, we examined differential expression of CR genes and the R6 co-expressing signatures (R6R3, R6R5, R6R3R5) across ICB responder/non-responder clinical groups using average gene expression pseudobulked by sample (summation of counts in each sample divided by total number of cells in each sample) to account for CR expression sparsity and intrapatient heterogeneity. Pseudobulked sample gene expression and signature expression were compared across clinical groups using boxplots and significance was assessed using Mann-Whitney U-tests. Code relating to this data analysis is publicly available on GitHub: https://github.com/azizilab/chemokine_receptor_reproducibility.

### Statistical analysis

One-way ANOVA was used for normally distributed data. Mann–Whitney was used for non-parametric data. Kaplan–Meier log-rank analysis was used to evaluate the survival differences between groups. Statistical analysis was conducted using Prism 10.0 software (GraphPad). Only p values < 0.05 were considered as significant.

---

**Extended Data Fig. 1.**
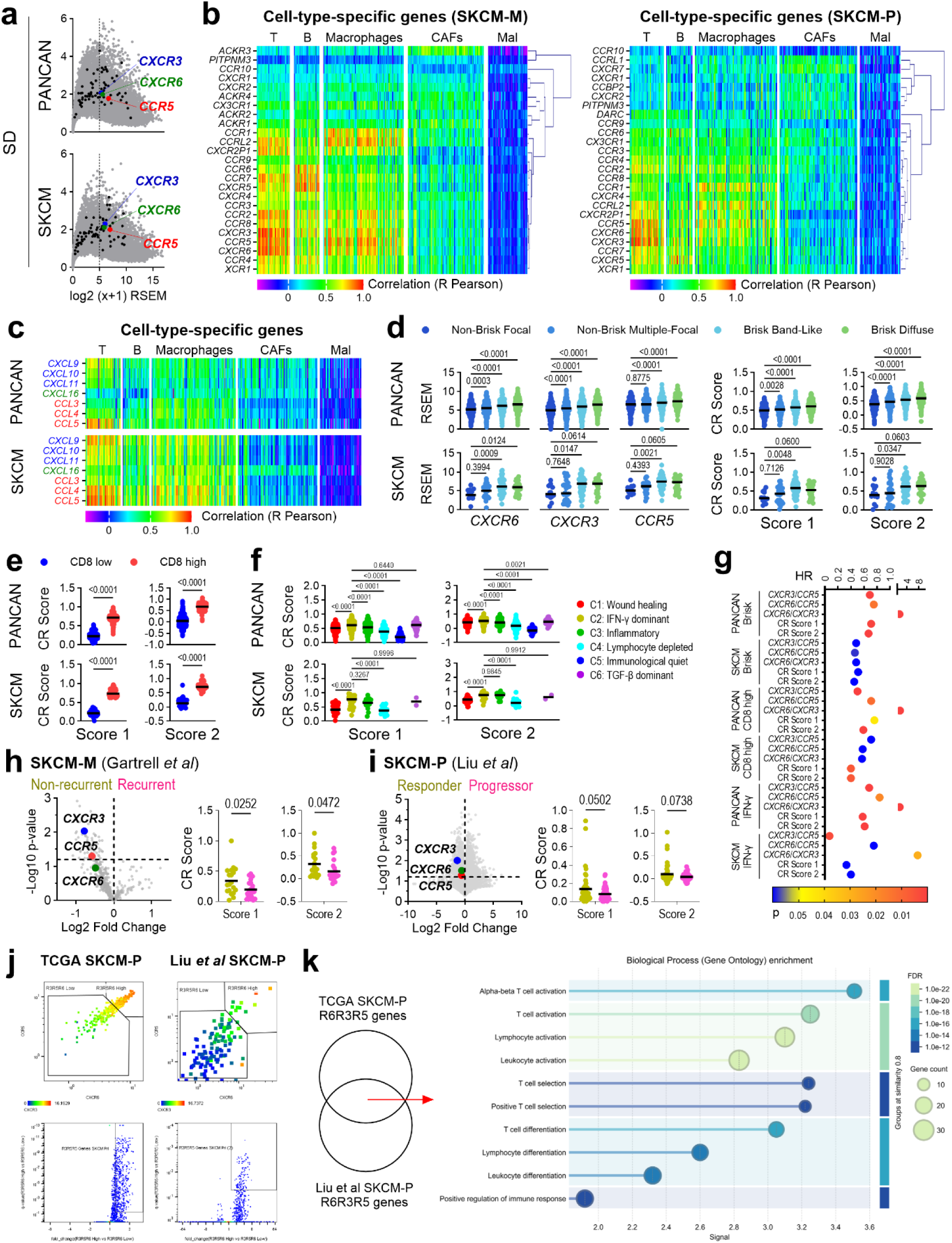
Gene expression of chemokine receptors and their ligands in human cancer. *In-silico* analysis of tumor transcriptome from TCGA Pan-Cancer Atlas (PANCAN; n=11323) and melanoma (SKCM; n=475) datasets. **a.** Mean-Variance Plot of gene expression, highlighting *CXCR3*, *CCR5* and *CXCR6*. Black dots represent genes for other CRs and chemokines. **b.** Hierarchical clustering heatmap of CR gene expression correlated with cell-type-specific gene signatures in metastatic (SKCM-M) and primary (SKCM-P) melanoma. **c**. Hierarchical clustering heatmap showing correlation of CXCR3, CCR5 and CXCR6 ligands gene expression with cell types in the tumor microenvironment. **d.** Expression of individual *CXCR6*, *CXCR3, CCR5,* and combinatorial CR scores (based on CXCR6, CXCR3 and CCR5 expression) across “brisk” classifications. **e, f**. CR scores stratified by “CD8^+^ score” (**e**) or immune subtype (**f**). **g**. Hazard ratios (HR) for survival prediction by CR scores and CR ratios specified tumor subtypes. **h, i**. CR scores in non-recurrent vs recurrent metastatic tumors (**h**) and responders vs progressors primary tumors (**i**). **j**. Upregulated differentially expressed genes (DEGs) in samples with high *CXCR6*, *CXCR3*, and *CCR5* co-expression. **k**. Gene Ontology analysis of shared upregulated DEGs.

**Extended Data Fig. 2.**
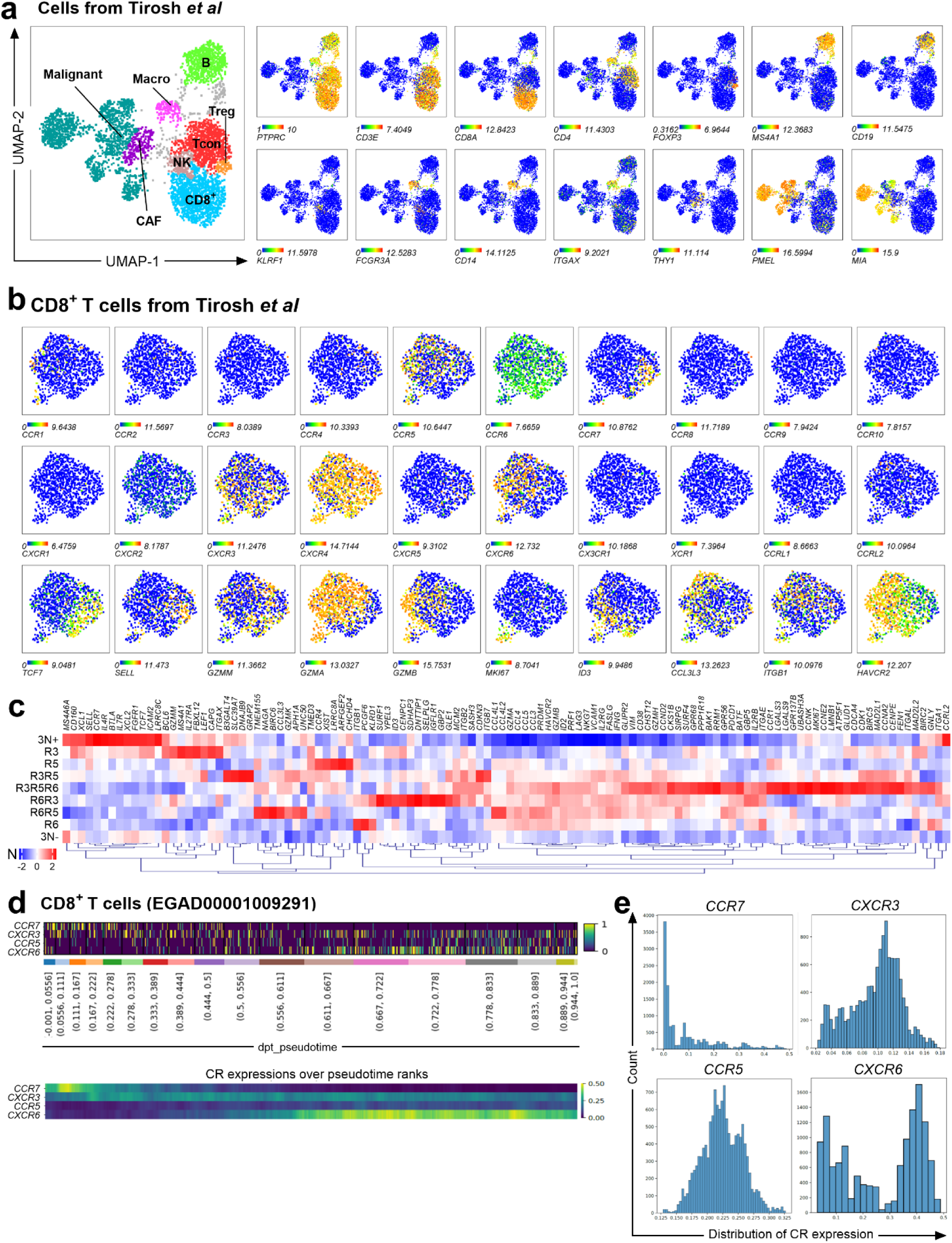
a, c. Single-cell Gene expression of chemokine receptors in human metastatic melanoma. **a.** UMAP projection of immune populations in the tumor microenvironment (GSE72056). **b.** Feature plots showing expression of naive, cytotoxicity, and exhaustion markers, transcription factors and CRs in CD8^+^ T cells. **c**. Hierarchical clustering heatmap of functional markers across CR-defined CD8+ T-cells. **d, e.** Trajectory analysis of CR expression in melanoma (EGAD00001009291). **d**. Rank sort and sliding window analysis (window size = 300 cells) of *CCR7*, *CXCR3*, *CCR5* and *CXCR6* across pseudotime. **e.** Distributions of normalized CR expression among CD8^+^ T cells.

**Extended Data Fig. 3.**
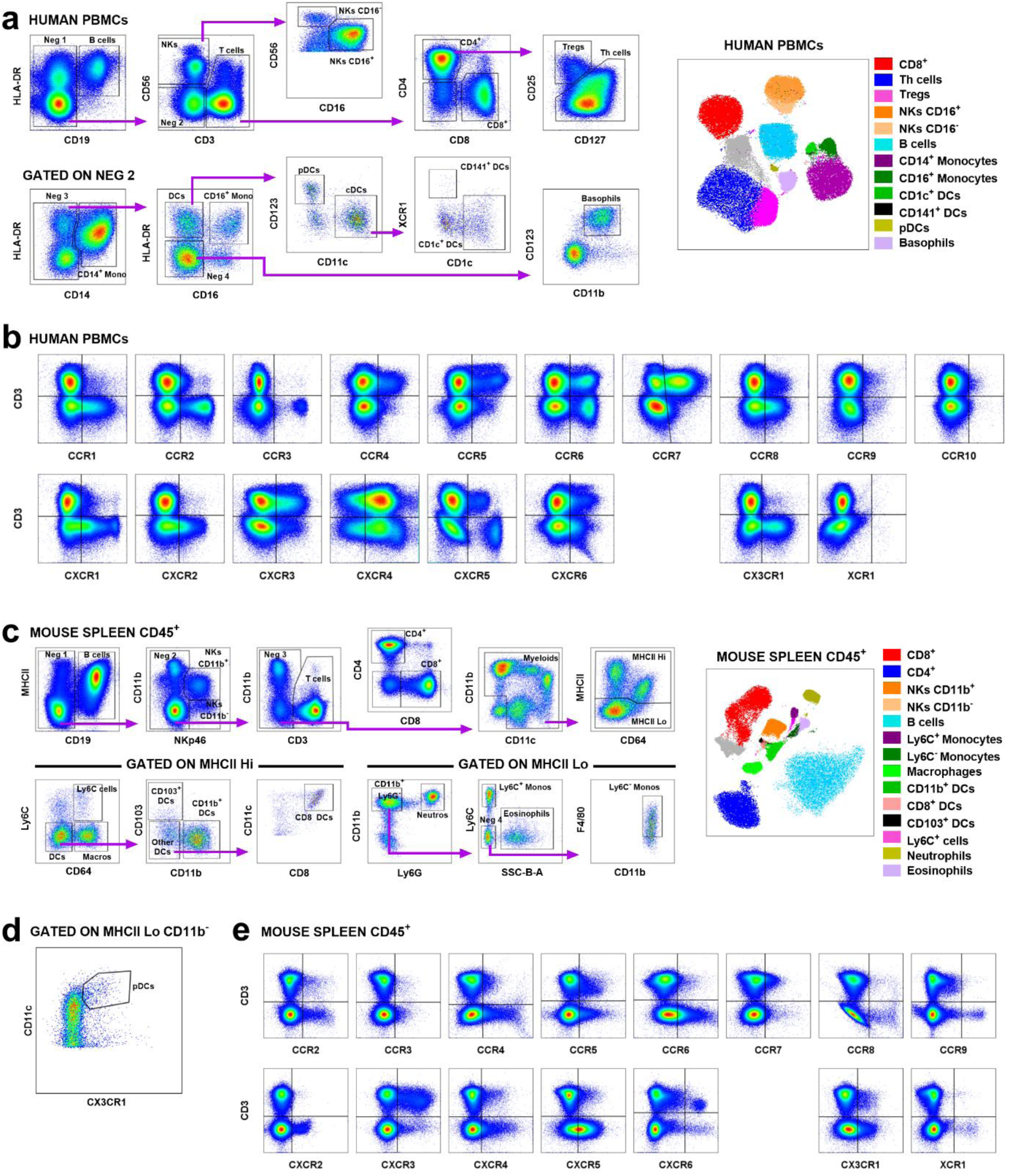
Gating strategy for spectral flow cytometry. Human PBMCs (n=11) and murine splenocytes (n=5) analyzed using a 43-color extracellular panel and 42-color intracellular panel (human) and a 36-color extracellular panel and a 35-color intracellular panel (mouse). **a.** Gating strategy and UMAP visualization for human immune subsets. **b.** Representative plots of CR expression in human PBMCs. **c, d.** Gating strategy for murine immune subsets. **e.** Representative plots of CR expression in murine splenocytes.

**Extended Data Fig. 4.**
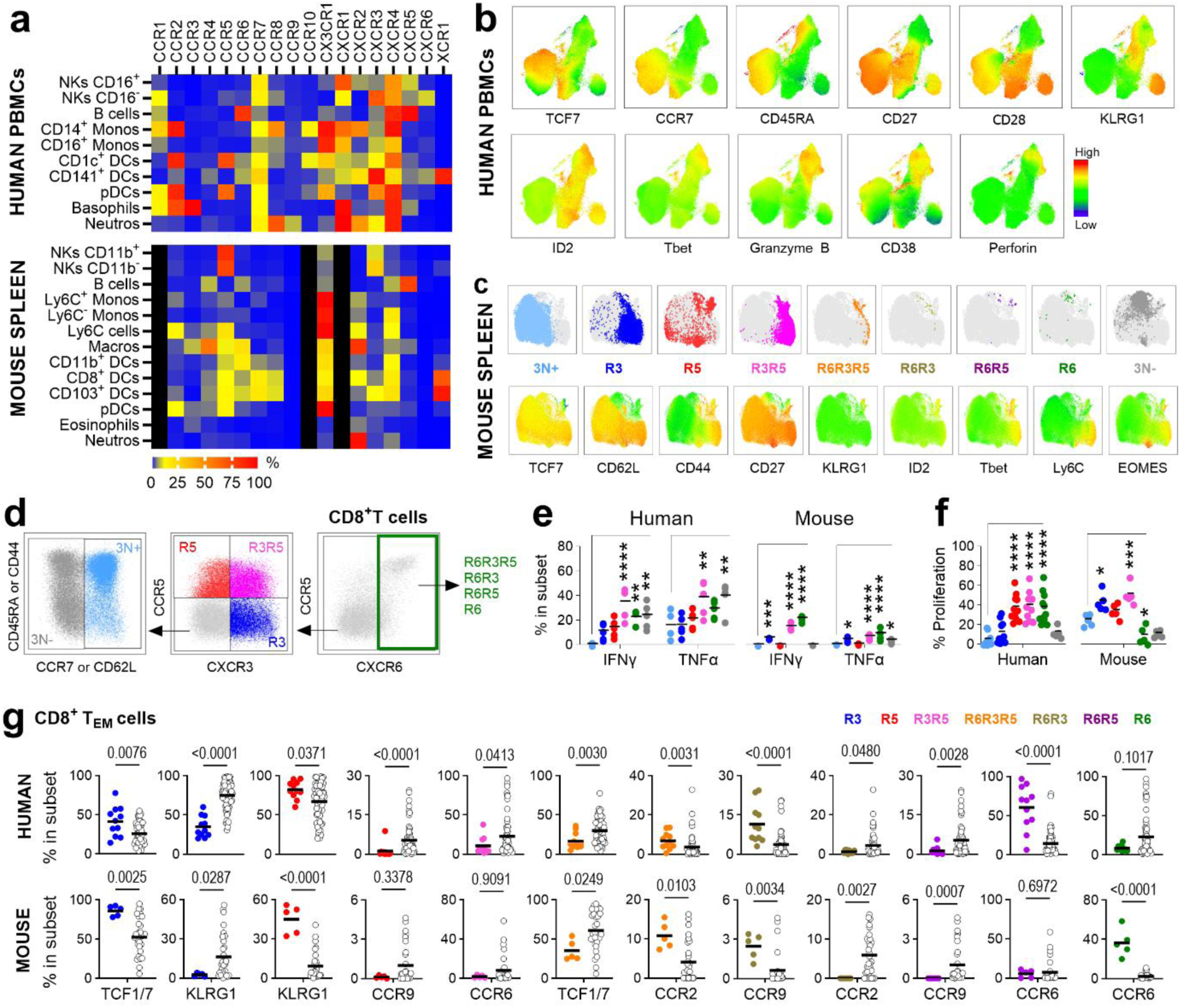
Functional and differentiation profiles of CR-defined subsets. **a**. Heatmap of CR heterogeneity across immune subsets in humans and mice. Grey indicates no data. **b**. UMAP projection showing functional markers in human CD8^+^ T cells. **c**. UMAP projection of CR-defined subsets overlaid with functional markers. **d-f**. Functional characterization of sorted and stimulated CR-defined subsets. **d**. Gating strategy for sorting. **e, f**. Activation (**e**) and proliferation (**f**) profiles of sorted subsets. **g**. Differential protein expression between specific CR subsets and all other subsets (white dots). Statistical significance calculated using unpaired ANOVA or Mann-Whitney multiple comparisons tests, *P < 0.05, **P < 0.01, ***P < 0.001, ****P < 0.0001.

**Extended Data Fig. 5.**
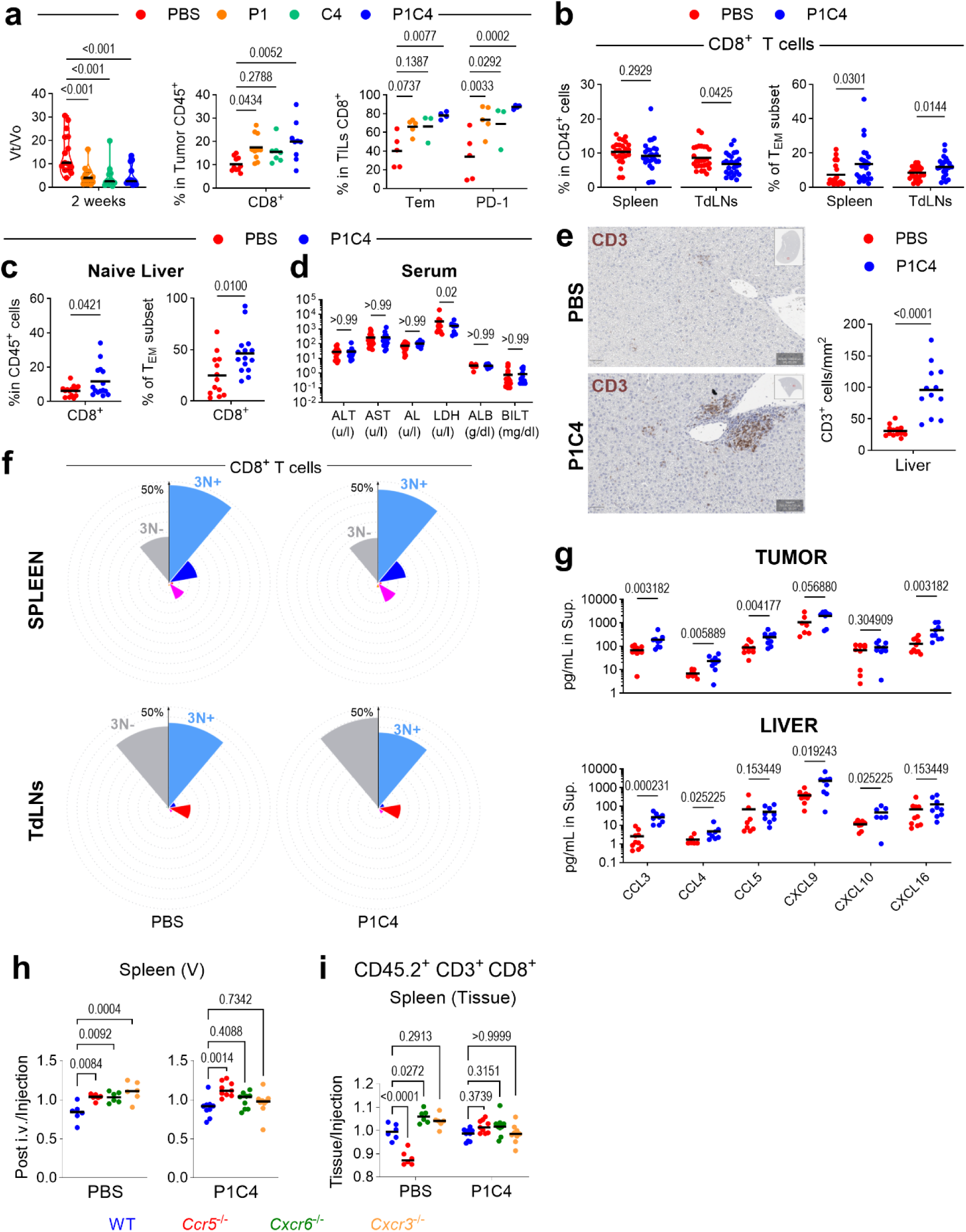
Dual PD-1/CTLA4 blockade increases T_EM_ cells in tumors and target tissues of Immune-Related Adverse Events. **a, b.** Analysis of B16-F10 tumor-bearing mice (n=5-15) were treated with monotherapy or combination (P1C4). **a**. Tumor volume (left, Vt = tumor volume at time t and Vo = volume at baseline) and CD8^+^ and T_EM_ (center/right). **b**. Treatment effect on percentages of CD8^+^ T cells and T_EM_ in spleen and TdLNs. **c**. Livers CD8^+^ T_EM_ infiltration in non-tumor bearing mice after treatment. **d**. Serum liver toxicity markers after treatment. **e**. Immunochemistry of CD3^+^ T-cells after treatment. **f**. Nightingale Rose of CR-defined subsets in spleen and TdLNs. **g**. Concentrations of CR ligands in tumor and liver supernatants. **h, i**. Competitive homing assay (described in Fig. 3i) controls showing pre-injection CD8^+^ T proportions (**h**) with a vascular increase (CD90.2^+^) of KO cells after 48 hours and splenic tissue/injection ratios (**i**). Statistical significance was calculated using unpaired ANOVA or Mann-Whitney multiple comparisons tests.

**Extended Data Fig. 6.**
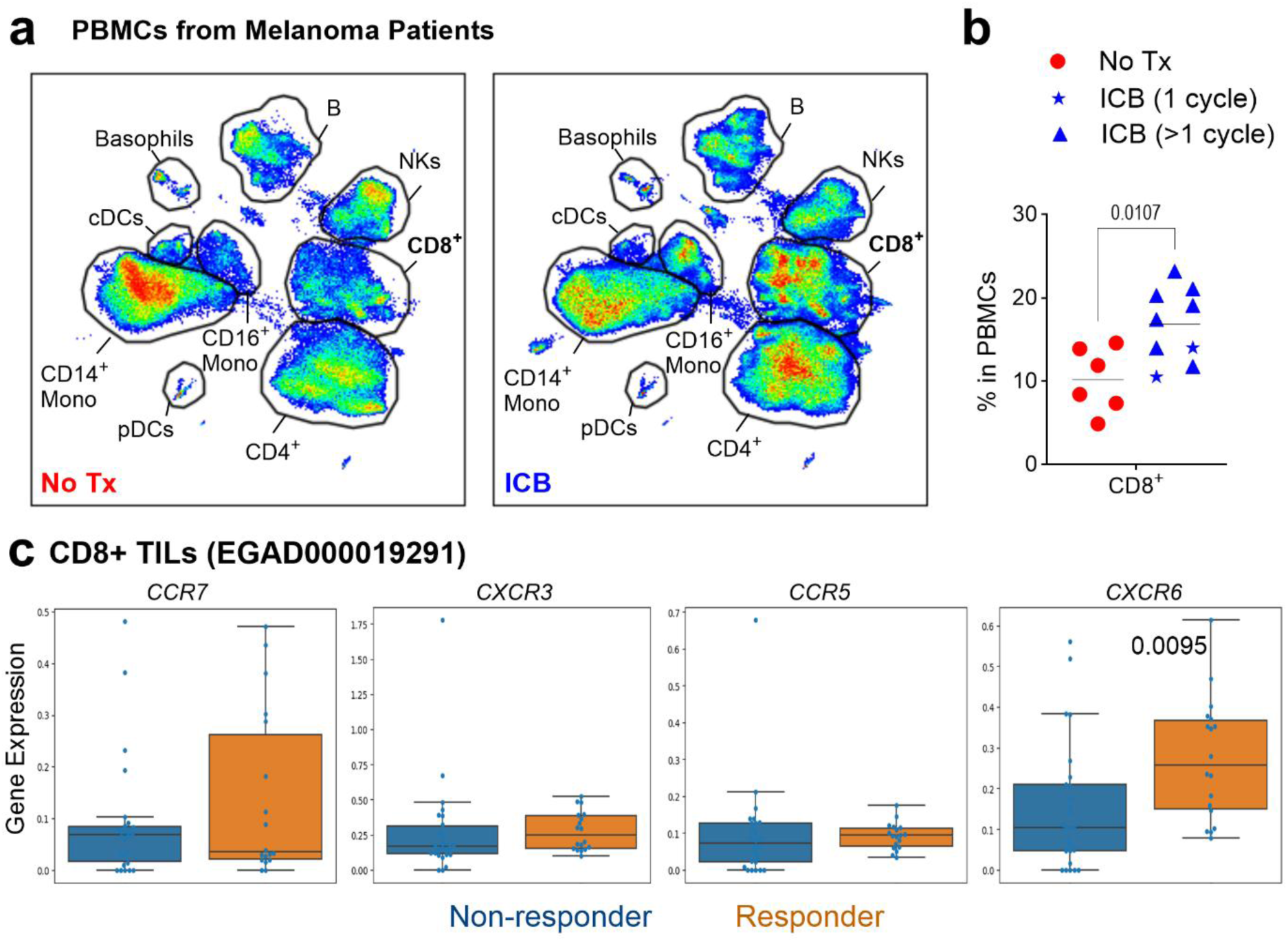
Peripheral T cell dynamics during ICB in melanoma patients. **a, b.** UMAP projection (**a**) and quantification (**b**) of CD8 T^+^ cells in PBMCs of melanoma patients. **c**. CR gene expression in ICB responders versus non-responders (EGAD000019291). Statistical significance determined by unpaired ANOVA or Mann-Whitney multiple comparisons tests.

## References

1 Wolchok, J. D. et al. Final, 10-Year Outcomes with Nivolumab plus Ipilimumab in Advanced Melanoma. N Engl J Med (2024). 10.1056/NEJMoa2407417

2 Morad, G., Helmink, B. A., Sharma, P. & Wargo, J. A. Hallmarks of response, resistance, and toxicity to immune checkpoint blockade. Cell 184, 5309–5337 (2021). 10.1016/j.cell.2021.09.020

3 Doki, Y. et al. Nivolumab Combination Therapy in Advanced Esophageal Squamous-Cell Carcinoma. N Engl J Med 386, 449–462 (2022). 10.1056/NEJMoa2111380

4 Wolchok, J. D. et al. Overall Survival with Combined Nivolumab and Ipilimumab in Advanced Melanoma. N Engl J Med 377, 1345–1356 (2017). 10.1056/NEJMoa1709684

5 Sznol, M. et al. Pooled Analysis Safety Profile of Nivolumab and Ipilimumab Combination Therapy in Patients With Advanced Melanoma. J Clin Oncol 35, 3815–3822 (2017). 10.1200/JCO.2016.72.1167

6 Motzer, R. J. et al. Nivolumab plus Ipilimumab versus Sunitinib in Advanced Renal-Cell Carcinoma. N Engl J Med (2018). 10.1056/NEJMoa1712126

7 Sharma, P. et al. Nivolumab Plus Ipilimumab for Metastatic Castration-Resistant Prostate Cancer: Preliminary Analysis of Patients in the CheckMate 650 Trial. Cancer Cell 38, 489–499 e483 (2020). 10.1016/j.ccell.2020.08.007

8 Hellmann, M. D. et al. Nivolumab plus Ipilimumab in Lung Cancer with a High Tumor Mutational Burden. N Engl J Med 378, 2093–2104 (2018). 10.1056/NEJMoa1801946

9 Overman, M. J. et al. Durable Clinical Benefit With Nivolumab Plus Ipilimumab in DNA Mismatch Repair-Deficient/Microsatellite Instability-High Metastatic Colorectal Cancer. J Clin Oncol, JCO2017769901 (2018). 10.1200/JCO.2017.76.9901

10 Gao, J. et al. Neoadjuvant PD-L1 plus CTLA-4 blockade in patients with cisplatin-ineligible operable high-risk urothelial carcinoma. Nat Med 26, 1845–1851 (2020). 10.1038/s41591-020-1086-y

11 Scherpereel, A. et al. Nivolumab or nivolumab plus ipilimumab in patients with relapsed malignant pleural mesothelioma (IFCT-1501 MAPS2): a multicentre, open-label, randomised, non-comparative, phase 2 trial. Lancet Oncol 20, 239–253 (2019). 10.1016/S1470-2045(18)30765-4

12 Johncilla, M. et al. Ipilimumab-associated Hepatitis: Clinicopathologic Characterization in a Series of 11 Cases. Am J Surg Pathol 39, 1075–1084 (2015). 10.1097/PAS.0000000000000453

13 Chen, P. L. et al. Analysis of Immune Signatures in Longitudinal Tumor Samples Yields Insight into Biomarkers of Response and Mechanisms of Resistance to Immune Checkpoint Blockade. Cancer Discov 6, 827–837 (2016). 10.1158/2159-8290.CD-15-1545

14 Daud, A. I. et al. Tumor immune profiling predicts response to anti-PD-1 therapy in human melanoma. J Clin Invest 126, 3447–3452 (2016). 10.1172/JCI87324

15 Rosenberg, S. A. et al. Durable complete responses in heavily pretreated patients with metastatic melanoma using T-cell transfer immunotherapy. Clin Cancer Res 17, 4550–4557 (2011). 10.1158/1078-0432.CCR-11-0116

16 Rohaan, M. W. et al. Tumor-Infiltrating Lymphocyte Therapy or Ipilimumab in Advanced Melanoma. N Engl J Med 387, 2113–2125 (2022). 10.1056/NEJMoa2210233

17 Sarnaik, A. A. et al. Lifileucel, a Tumor-Infiltrating Lymphocyte Therapy, in Metastatic Melanoma. J Clin Oncol 39, 2656–2666 (2021). 10.1200/JCO.21.00612

18 Schulz, O., Hammerschmidt, S. I., Moschovakis, G. L. & Forster, R. Chemokines and Chemokine Receptors in Lymphoid Tissue Dynamics. Annu Rev Immunol 34, 203–242 (2016). 10.1146/annurev-immunol-041015-055649

19 Bedognetti, D. et al. CXCR3/CCR5 pathways in metastatic melanoma patients treated with adoptive therapy and interleukin-2. Br J Cancer 109, 2412–2423 (2013). 10.1038/bjc.2013.557

20 Chheda, Z. S., Sharma, R. K., Jala, V. R., Luster, A. D. & Haribabu, B. Chemoattractant Receptors BLT1 and CXCR3 Regulate Antitumor Immunity by Facilitating CD8+ T Cell Migration into Tumors. J Immunol 197, 2016–2026 (2016). 10.4049/jimmunol.1502376

21 Hoch, T. et al. Multiplexed imaging mass cytometry of the chemokine milieus in melanoma characterizes features of the response to immunotherapy. Sci Immunol 7, eabk1692 (2022). 10.1126/sciimmunol.abk1692

22 Bell, B., Flores-Lovon, K., Cueva-Chicana, L. A. & Macedo, R. Role of chemokine receptors in gastrointestinal mucosa. Int Rev Cell Mol Biol 388, 20–52 (2024). 10.1016/bs.ircmb.2024.02.003

23 Valdivia-Silva, J. & Chinney-Herrera, A. Chemokine receptors and their ligands in breast cancer: The key roles in progression and metastasis. Int Rev Cell Mol Biol 388, 124–161 (2024). 10.1016/bs.ircmb.2024.07.002

24 Tang, Y., Hu, Y., Niu, Y., Sun, L. & Guo, L. CCL5 as a Prognostic Marker for Survival and an Indicator for Immune Checkpoint Therapies in Small Cell Lung Cancer. Front Med (Lausanne) 9, 834725 (2022). 10.3389/fmed.2022.834725

25 Seitz, S. et al. CXCL9 inhibits tumour growth and drives anti-PD-L1 therapy in ovarian cancer. Br J Cancer 126, 1470–1480 (2022). 10.1038/s41416-022-01763-0

26 Han, Y. et al. CXCL10 and CCL5 as feasible biomarkers for immunotherapy of homologous recombination deficient ovarian cancer. Am J Cancer Res 13, 1904–1922 (2023).

27 Reschke, R. et al. Immune cell and tumor cell-derived CXCL10 is indicative of immunotherapy response in metastatic melanoma. J Immunother Cancer 9 (2021). 10.1136/jitc-2021-003521

28 Fei, L., Ren, X., Yu, H. & Zhan, Y. Targeting the CCL2/CCR2 Axis in Cancer Immunotherapy: One Stone, Three Birds? Front Immunol 12, 771210 (2021). 10.3389/fimmu.2021.771210

29 Velasco-Velazquez, M. et al. CCR5 antagonist blocks metastasis of basal breast cancer cells. Cancer research 72, 3839–3850 (2012). 10.1158/0008-5472.CAN-11-3917

30 Sicoli, D. et al. CCR5 receptor antagonists block metastasis to bone of v-Src oncogene-transformed metastatic prostate cancer cell lines. Cancer research 74, 7103–7114 (2014). 10.1158/0008-5472.CAN-14-0612

31 Halama, N. et al. Tumoral Immune Cell Exploitation in Colorectal Cancer Metastases Can Be Targeted Effectively by Anti-CCR5 Therapy in Cancer Patients. Cancer Cell 29, 587–601 (2016). 10.1016/j.ccell.2016.03.005

32 Bachelerie, F. et al. International Union of Basic and Clinical Pharmacology. [corrected]. LXXXIX. Update on the extended family of chemokine receptors and introducing a new nomenclature for atypical chemokine receptors. Pharmacol Rev 66, 1–79 (2014). 10.1124/pr.113.007724

33 Saltz, J., Gupta, R., Hou, L., Kurc, T., Singh, P., Nguyen, V., Samaras, D., Shroyer, K. R., Zhao, T., Batiste, R., Van Arnam, J., The Cancer Genome Atlas Research Network, Shmulevich, I., Rao, A. U. K., Lazar, A. J., Sharma, A., & Thorsson, V. Tumor-Infiltrating Lymphocytes Maps from TCGA H&E Whole Slide Pathology Images [Data set]. (2018). 10.7937/K9/TCIA.2018.Y75F9W1

34 Wolf, D. Immune Signature Scores [Data set, https://pancanatlas.xenahubs.net]. (2016).

35 Thorsson, V. et al. The Immune Landscape of Cancer. Immunity 51, 411–412 (2019). 10.1016/j.immuni.2019.08.004

36 Gartrell, R. D. et al. Validation of Melanoma Immune Profile (MIP), a Prognostic Immune Gene Prediction Score for Stage II-III Melanoma. Clin Cancer Res 25, 2494–2502 (2019). 10.1158/1078-0432.CCR-18-2847

37 Liu, D. et al. Integrative molecular and clinical modeling of clinical outcomes to PD1 blockade in patients with metastatic melanoma. Nat Med 25, 1916–1927 (2019). 10.1038/s41591-019-0654-5

38 Tirosh, I. et al. Dissecting the multicellular ecosystem of metastatic melanoma by single-cell RNA-seq. Science 352, 189–196 (2016). 10.1126/science.aad0501

39 Pozniak, J. et al. A TCF4-dependent gene regulatory network confers resistance to immunotherapy in melanoma. Cell 187, 166–183 e125 (2024). 10.1016/j.cell.2023.11.037

40 Saelens, W., Cannoodt, R., Todorov, H. & Saeys, Y. A comparison of single-cell trajectory inference methods. Nat Biotechnol 37, 547–554 (2019). 10.1038/s41587-019-0071-9

41 Yanagihara, T., Tanaka, K. & Matsumoto, K. A measuring method for occupancy of immune checkpoint inhibitors in the cell surface. Biochem Biophys Res Commun 527, 213–217 (2020). 10.1016/j.bbrc.2020.04.122

42 Wong, M. T. et al. A High-Dimensional Atlas of Human T Cell Diversity Reveals Tissue-Specific Trafficking and Cytokine Signatures. Immunity 45, 442–456 (2016). 10.1016/j.immuni.2016.07.007

43 Berahovich, R. D., Lai, N. L., Wei, Z., Lanier, L. L. & Schall, T. J. Evidence for NK cell subsets based on chemokine receptor expression. J Immunol 177, 7833–7840 (2006). 10.4049/jimmunol.177.11.7833

44 Franciszkiewicz, K., Boissonnas, A., Boutet, M., Combadiere, C. & Mami-Chouaib, F. Role of chemokines and chemokine receptors in shaping the effector phase of the antitumor immune response. Cancer Res 72, 6325–6332 (2012). 10.1158/0008-5472.CAN-12-2027

45 Ozga, A. J., Chow, M. T. & Luster, A. D. Chemokines and the immune response to cancer. Immunity 54, 859–874 (2021). 10.1016/j.immuni.2021.01.012

46 Haghverdi, L., Buettner, F. & Theis, F. J. Diffusion maps for high-dimensional single-cell analysis of differentiation data. Bioinformatics 31, 2989–2998 (2015). 10.1093/bioinformatics/btv325

47 Villablanca, E. J. & Mora, J. R. Competitive homing assays to study gut-tropic t cell migration. J Vis Exp (2011). 10.3791/2619

48 Anderson, K. G. et al. Intravascular staining for discrimination of vascular and tissue leukocytes. Nat Protoc 9, 209–222 (2014). 10.1038/nprot.2014.005

49 Aran, D. et al. Reference-based analysis of lung single-cell sequencing reveals a transitional profibrotic macrophage. Nat Immunol 20, 163–172 (2019). 10.1038/s41590-018-0276-y

50 Lopez, R., Regier, J., Cole, M. B., Jordan, M. I. & Yosef, N. Deep generative modeling for single-cell transcriptomics. Nat Methods 15, 1053–1058 (2018). 10.1038/s41592-018-0229-2

## Methods Bibliography

51. Goldman, M.J., Craft, B., Hastie, M. et al. Visualizing and interpreting cancer genomics data via the Xena platform. Nat Biotechnol (2020). 10.1038/s41587-020-0546-8

52. Saeed AI, Sharov V, White J, Li J, Liang W, Bhagabati N, Braisted J, Klapa M, Currier T, Thiagarajan M, Sturn A, Snuffin M, Rezantsev A, Popov D, Ryltsov A, Kostukovich E, Borisovsky I, Liu Z, Vinsavich A, Trush V, Quackenbush J. TM4: a free, open-source system for microarray data management and analysis. Biotechniques. 2003 Feb;34(2):374–8

53. Lanczky A, Gyorffy B. Web-Based Survival Analysis Tool Tailored for Medical Research (KMplot): Development and Implementation, J Med Internet Res, 2021; 23(7):e27633, doi:10.2196/27633

54. Szklarczyk D, Kirsch R, Koutrouli M, Nastou K, Mehryary F, Hachilif R, Gable AL, Fang T, Doncheva NT, Pyysalo S, Bork P, Jensen LJ, von Mering C. The STRING database in 2023: protein-protein association networks and functional enrichment analyses for any sequenced genome of interest. Nucleic Acids Res. 2023 Jan 6;51(D1):D638–D646. doi: 10.1093/nar/gkac1000.

