## Supplementary Figures for "A Conserved Landscape of Chemokine Receptor Co-expression Defines the Functional States of CD8+ T Cells in Melanoma"

**Supplementary Figure 1.**

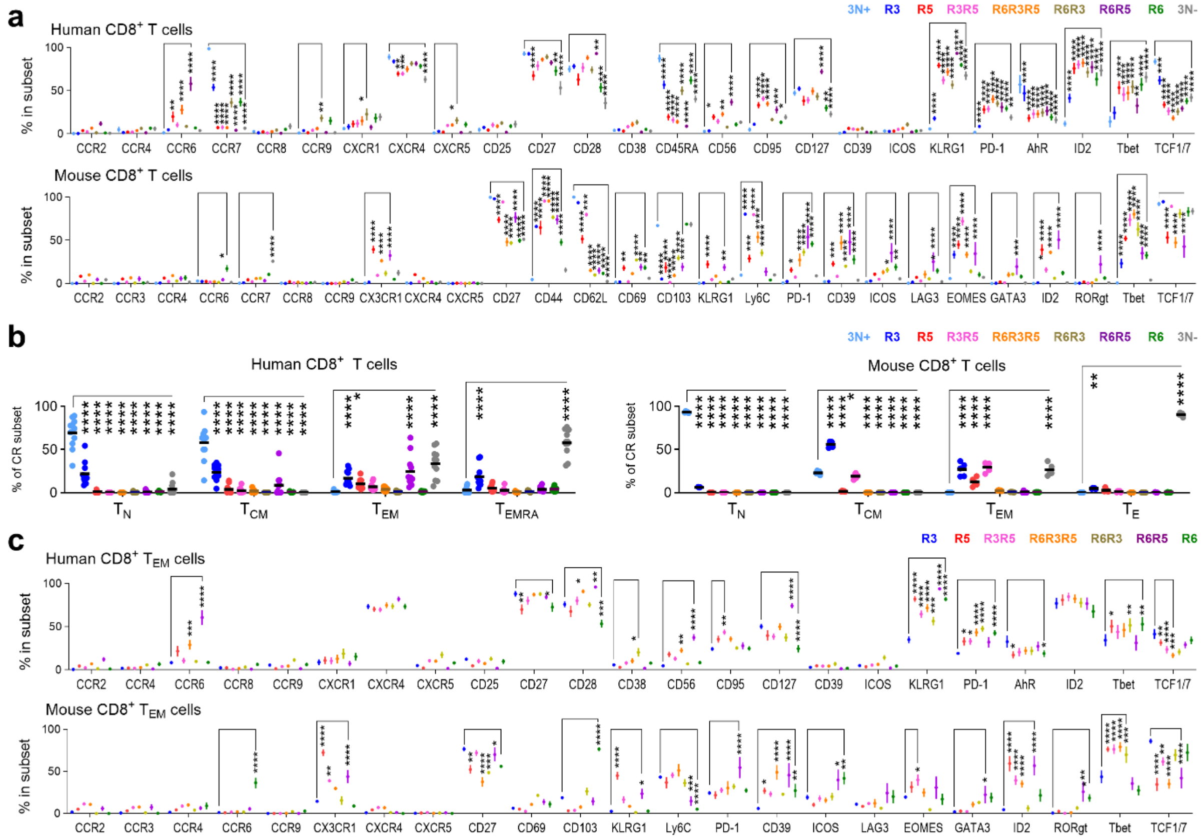


**Supplementary Figure 1. Phenotypic and functional characterization of CR-defined T cell subsets. a.** Expression of CRs, transcription factors, activation and exhaustion markers across CR-defined $CD8^+$ T-cell subsets (corresponding to Fig. 2d). **b**. Distribution of CR-defined subsets within T-cell memory subsets (corresponding to Fig. 2e). **c**. Expression of CRs, transcription factors, activation and exhaustion markers among CR-defined $CD8^+$ $T_{EM}$ cell subsets (corresponding to Fig. 2f). Statistical significance determined by unpaired Mann-Whitney tests, *P < 0.05, **P < 0.01, ***P < 0.001, ****P < 0.0001.

**Supplementary Figure 2.**

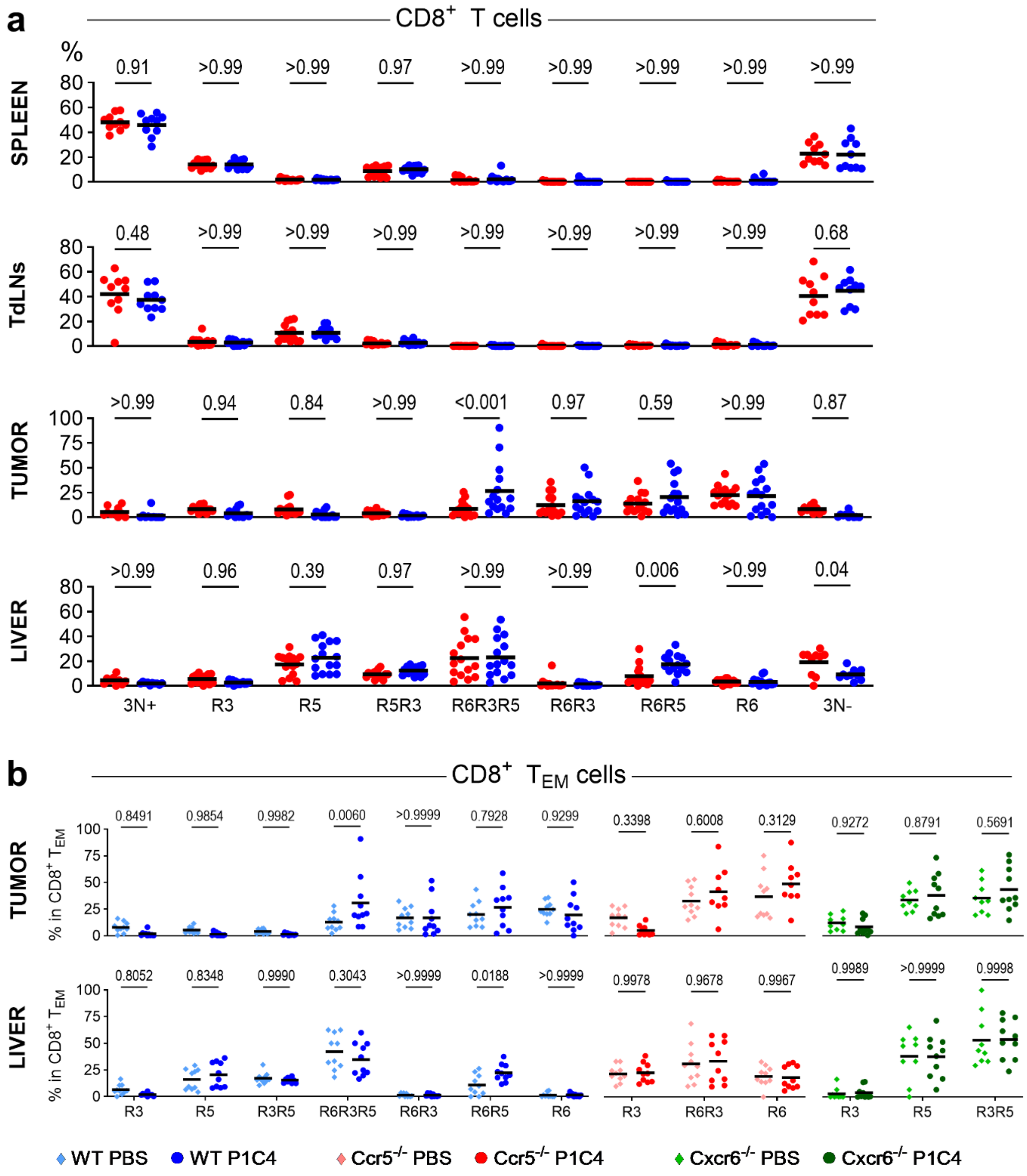


**Supplementary Figure 2. Tissue-specific modulation of CR-defined subsets by immune checkpoint blockade. a.** Percentages of CR-defined subsets in spleen, tumor-draining lymph nodes (TdLNs), tumor and liver following P1C4 treatment compared to PBS control group (expanded data for Fig. 3c and Extended Data Fig. 5f). **b**. Genetic deletion of *Ccr5* or *Cxcr6* abrogates the treatment-induced expansion of these subsets (expanded data for Fig. 3e. Statistical significance determined by unpaired Mann-Whitney tests.

**Supplementary Figure 3.**

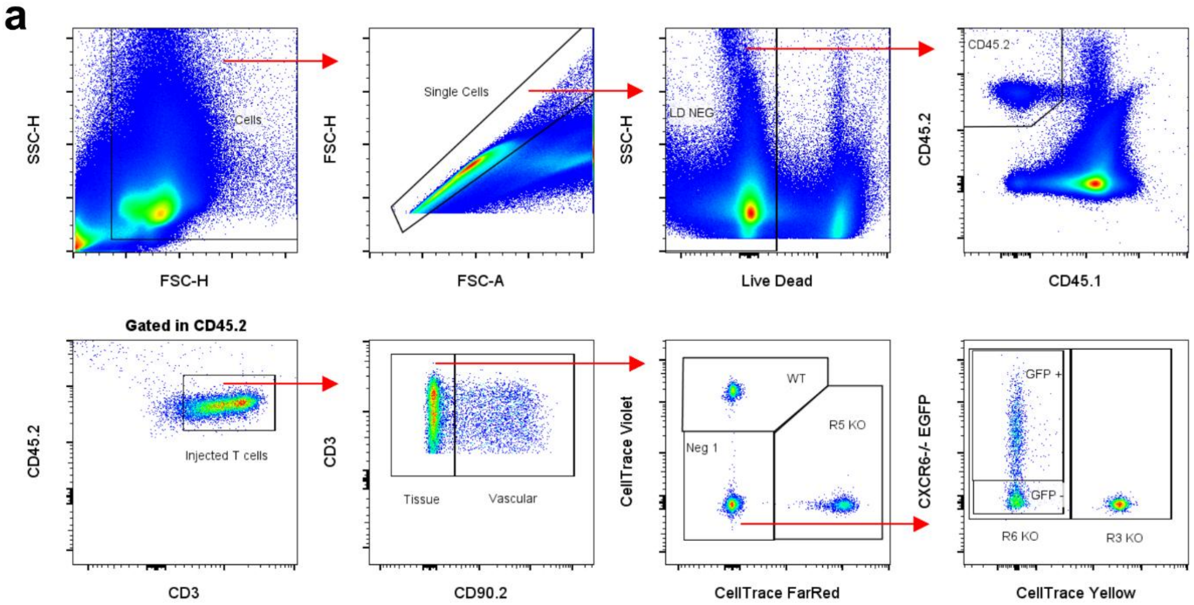


**Supplementary Figure 3. Gating strategy for competitive homing assay. a.** Flow cytometry gating strategy used to identify and track adoptively transferred T cells in recipient tissues 48 hours post-injection.

**Supplementary Figure 4.**

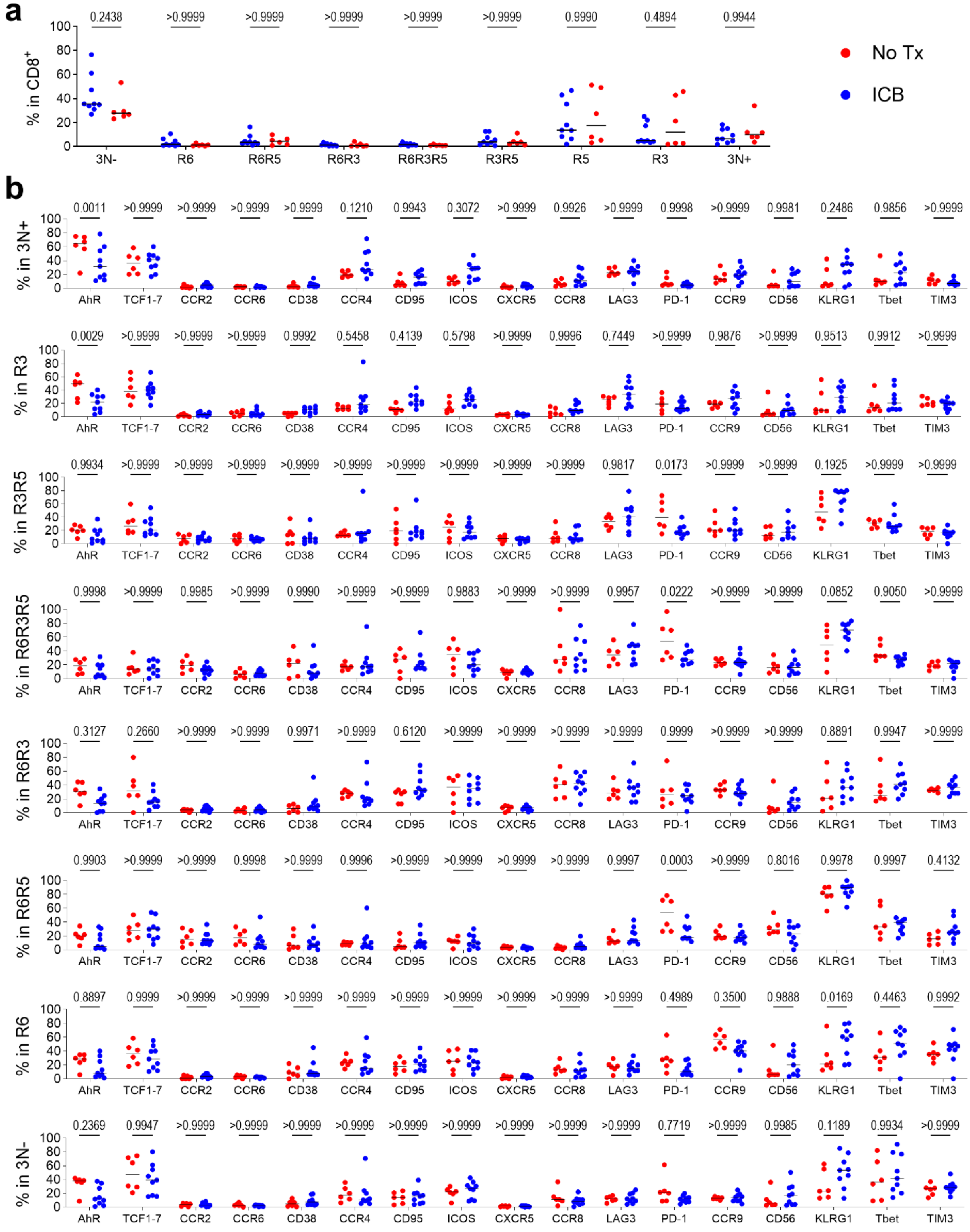

**Supplementary Figure 4. Systemic impact of ICB on CR-defined subset frequency and phenotype. a.** Percentages of CR-defined CD8$^+$ T-cell subsets in PBMCs from ICB-treated melanoma patients versus untreated (No Tx) controls (expanded data for Fig. 4b right). **b**. Expression of CRs, transcription factors, activation and exhaustion markers in circulating CR-defined CD8$^+$ T-cell subsets following ICB treatment (expanded data for Fig. 4c). Statistical significance determined by unpaired Mann-Whitney tests.
